# Inactivating a β1 subunit of SnRK1 promotes broad-spectrum disease resistance in rice

**DOI:** 10.1101/2025.06.26.661071

**Authors:** Guixin Yuan, Xunli Lu, Xingbin Wang, Mengfei Li, Shiwei Wang, Zhaoxiang Huang, Fengrui Zhang, Xin Zhang, Jun Yang, Hailong Guo, Vijai Bhadauria, Wangsheng Zhu, Wensheng Zhao, You-Liang Peng

## Abstract

Rice is a staple crop for over half of the world’s population, and its sustainable production is vital to ensure global food security. However, rice is highly susceptible to several devastating fungal diseases^1^, including blast disease caused by *Magnaporthe oryzae*, sheath blight by *Rhizoctonia solani*, false smut by *Ustilaginoidea virens*, brown spot by *Bipolaris oryzae*, bakanae by *Fusarium fujikuroi*, and head blight by *F. graminearum*. The mechanisms underlying the susceptibility of rice to these fungal diseases remain unclear. Here, we report that the β subunit of SnRK1, SnRK1β1A, confers broad-spectrum susceptibility to these fungal diseases. Our findings reveal that diverse rice fungal pathogens have convergently evolved an effector, Gas2, which interacts with SnRK1β1A to prevent its ubiquitination-mediated degradation and promotes its nuclear translocation. While *SnRK1*β*1A* is expressed at low levels in healthy plants, its expression is markedly induced upon fungal infections, facilitating susceptibility by inhibiting SnRK1α1, the α subunit of SnRK1, whose overexpression has been shown to enhance broad-spectrum resistance in rice^2^. Notably, rice lines with disrupted *SnRK1*β*1A* are resistant to multiple fungal diseases without compromising growth and yield in the field. Together, this study demonstrates that broad-spectrum disease resistance in crops can be achieved by disrupting inducible susceptibility genes whose encoded proteins are targeted by effectors conserved in multiple pathogens.

## Main

Like all other species of plants, rice plants are attacked by a number of fungal diseases throughout their growth, development, and production stages^1^. Several of these diseases are particularly devastating, including blast disease, sheath blight, false smut, brown spot, bakanae and head blight, each of which can lead to significant yield losses, often exceeding 1% both regionally or globally^3^. Addressing these fungal disease threats presents a critical challenge for ensuring global food security. Among the available strategies, utilizing resistant cultivars remains the most economical and environmentally friendly measure for controlling plant diseases. However, no commercial rice cultivars currently on the market possess broad-spectrum resistance to these fungal diseases.

Susceptibility (*S*) genes are plant genes that pathogens co-opt to facilitate infections, and their loss-of-function typically enhances disease resistance^4-6^. Hundreds of *S* genes have been identified across various plant species, with recessive *S* alleles being successfully utilized in crop resistance breeding programs^5,6^, such as *mlo11*^7^ and *pi21*^8^. Notably, advancements in CRISPR/Cas9-mediated gene-editing technology have positioned *S* gene disruption as a promising strategy for engineering disease-resistant plants^9,10^. Successful examples include the precise editing of *Mlo* and *TaPsIPK1* in wheat^11,12^, *Pi21*, *RBL1*^Δ*12*^ and *SWEETs* in rice^13-16^, and *ZmNANMT* in maize^17^, *SIDMR6-1* in tomato^18^, and *StPM1* in potato^19^. However, beyond acting as susceptibility factors, most *S* genes play critical roles in plant growth and development, and their disruption can result in growth and yield penalties^16,20,21^. Therefore, the challenge remains to efficiently identify *S* genes whose inhibition can confer broad-spectrum disease resistance without compromising plant fitness and productivity.

### Multiple rice fungal pathogens possess a conserved *MAS* gene

Previous studies have reported that *Magnaporthe* appressorium-specific (MAS) proteins are widely distributed in fungi, with some playing essential roles in virulence^22-24^. We, therefore, wondered whether there exists a conserved MAS effector in rice fungal pathogens and then searched for MAS proteins in *M. oryzae, B. oryzae*, *F. fujikuroi*, *F. graminearum*, *R. solani*, and *U. virens*. The search revealed that these fungal pathogens possess eight, four, five, four, thirteen, and one MAS proteins, respectively (Extended Data Fig. 1a). Secondary structure analysis and the 3D structure prediction suggested that *M. oryzae, B. oryzae*, *R. solani*, *F. graminearum*, and *F. fujikuroi* each contain two, while *U. virens* has one homolog of Gas2, a known MAS protein in *M. oryzae* required for virulence^23,25^ (Extended Data Fig. 1a,b).

To determine whether these proteins are functional orthologs of Gas2, we performed gene complementation by independently introducing these Gas2 orthologs from the individual fungal pathogens into Δ*gas2*, a *GAS2* deletion mutant of *M. oryzae* (Extended Data Fig. 1c). The complementation showed that *XP_007693280.1* (*BoGAS2*) from *B. oryzae*, *CCO30280.1* (*RsGAS2*) from *R. solani*, *XP_023432757.1* (*FfGAS2*) from *F. fujikuroi*, *XP_011320808.1* (*FgGAS2*) from *F. graminearum*, and *XP_043001658.1* (*UvGAS2*) from *U. virens* restored the wild type (WT) virulence of Δ*gas2* (Fig. 1a), whereas *XP_007687900.1* (named as *BoGAS2L*) from *B. oryzae*, and *XP_960032.1* (*NcMAS1*) and *XP_962708.1* (*NcMAS2*) from saprophytic *Neurospora crassa* failed to rescue the virulence of Δ*gas2* (Fig. 1a), indicating that Gas2 is conserved in multiple rice fungal pathogens.

**Fig. 1.**
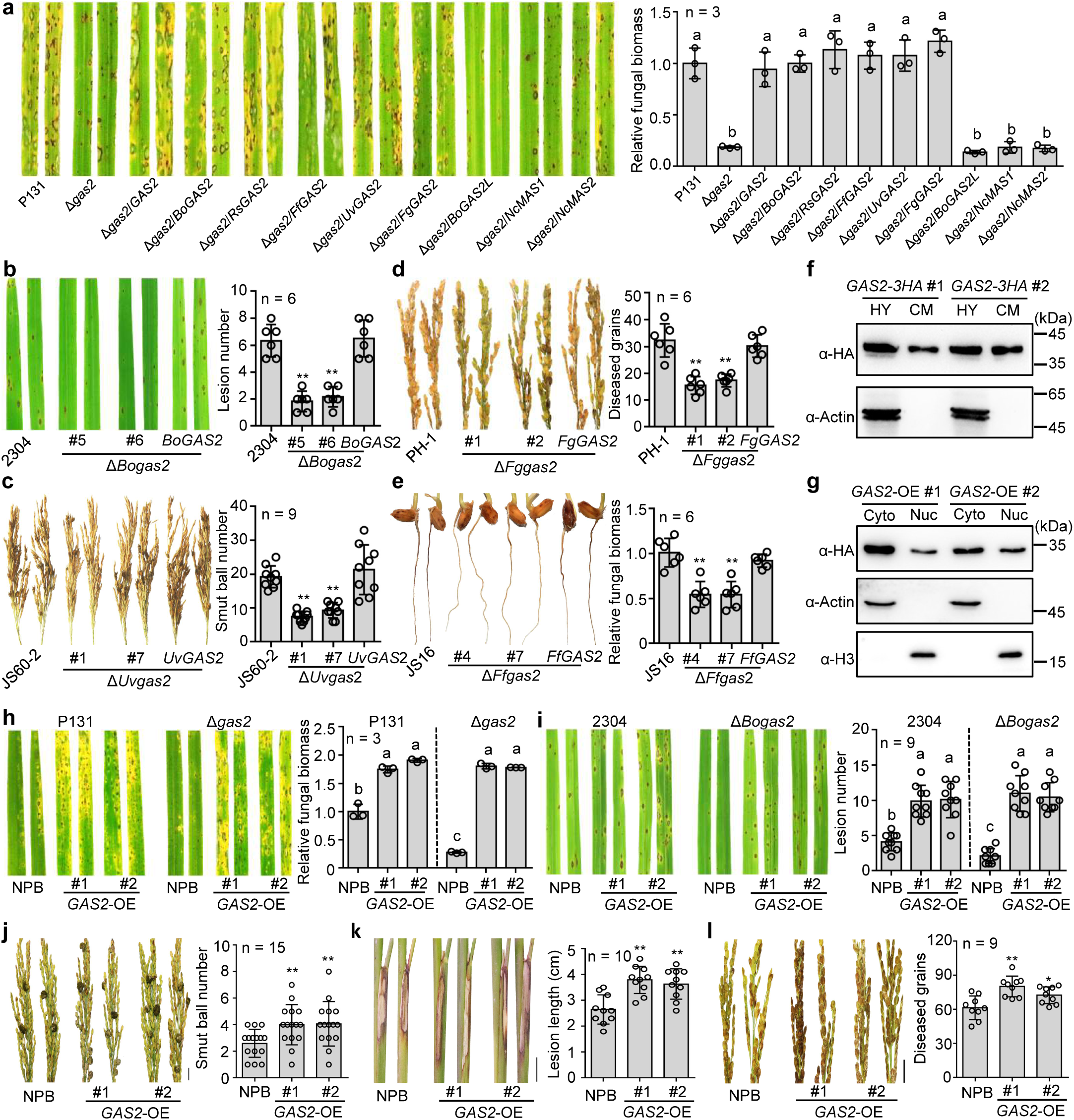
Gas2 is a virulence effector conserved in rice major fungal pathogens. **a,** Virulence of *M. oryzae* Δ*gas2* mutant on rice leaves is rescued by *GAS2*-orthologues but neither by *GAS2* paralogues (*BoGAS2L*) from *B. oryzae* nor by unrelated *MAS* from saprophytic *Neurospora crassa*. Relative fungal biomass in inoculated rice leaves was measured by qPCR at 5 days post-inoculation (dpi). **b-e,** *GAS2* is required for full virulence of *B. oryzae* (**b**), *U. virens* (**c**), *F. graminearum* (**d**) and *F. fujikuroi* (**e**). Disease severity was calculated for each pathogen. **f,** Gas2-3HA detected by western blotting (WB) with α-HA antibody in mycelium and culture medium of *M. oryzae* transformant. Actin is used as a loading control. **g,** 3HA-Gas2 detected by WB in the cytoplasmic and nuclear protein extracts of NPB lines expressing *GAS2* (*GAS2*-*OE*) using α-HA antibody. α-Actin and α-Histone H3 were used as cytoplasmic and nuclear markers, respectively. **h-l,** *GAS2*-*OE* lines showed heavier symptoms as compared with its wild type (WT) NPB after infection by *M. oryzae* strain P131 and its Δ*gas2* mutant (**h**), *B. oryzae* isolate 2304 and its Δ*Bogas2* mutant (**i**), *U. virens* isolate JS60-2 (**j**), *R. solani* isolate XN27 (**k**) and *F. graminearum* isolate PH-1 (**l**). Bar-graphs right to photos indicate disease severity caused by individual pathogens. Data represent mean ± SD. One-way ANOVA followed by post-hoc Turkey tests were used in (**a, h, i)**, letters indicate significantly different groups; and two-tailed Student’s t-test was used in (**b-e, j-l)**. *p < 0.05, **p < 0.01.

We further generated *GAS2* deletion mutants in *B. oryzae*, *F. fujikuroi*, *U. virens*, and *F. graminearum* (Extended Data Fig. 1d-g). Due to the lack of a genetic transformation system for *R. solani*, we were unable to generate deletion mutants in this species. All the gene deletion mutants in the respective fungal pathogens exhibited a dramatic reduction in virulence compared to their isogenic strains (Fig. 1b-e), verifying that *GAS2* orthologs are required for the virulence of the rice fungal pathogens. Additionally, complementation of Δ*Fggas2* (an *FgGAS2* deletion mutant of *F. graminearum* that causes *Fusarium* head blight in rice) revealed that *GAS2* from *M. oryzae*, *R. solani*, *F. fujikuroi*, *U. virens*, and *F. graminearum* restored the WT virulence of Δ*Fggas2* (Extended Data Fig. 1h), further confirming that *GAS2* is functionally conserved in rice fungal pathogens. All Gas2 proteins contain an α helix at the C termini (Extended Data Fig. 1a,b). Gene complementation of Δ*gas2* using *GAS2*^Δ*C*^, which encodes a truncated Gas2 lacking the α helix, showed that Δ*gas2/GAS2*^Δ*C*^ transformants displayed virulence similar to Δ*gas2* (Extended Data Fig. 1i), indicating that the C terminal α helix is essential to the virulence function of Gas2.

### Gas2 is an intracellular effector suppressing rice PTI

MAS proteins of *Podosphaera xanthii* have been reported to exhibit chitinase activity^25^. We expressed the Gas2 protein of *M. oryzae* in *Escherichia coli* but were unable to detect its chitinase activity (Extended Data Fig. 2a). To investigate the functional characteristics of Gas*2* of *M. oryzae*, we first evaluated its expression by RT-qPCR, revealing that *GAS2* was highly expressed during appressorium-mediated penetration and primary infection hyphae development but expressed at low levels during vegetative hyphae growth and necrotrophic development in plant tissues (Extended Data Fig. 2b). We further inoculated rice leaf sheaths with a Δ*gas2*/*GAS2-GFP* strain to check the Gas2-GFP expression profile. The GFP fluorescence signals were only observed in the appressorium from 4 hours post-inoculation (hpi) to 12 hpi (Extended Data Fig. 2c), confirming that Gas2 is mainly expressed during appressorium penetration.

We then tested whether Gas2 is a secreted protein. Gas2 has a predicted signal peptide (SP) of 22 amino acids at its N-terminus. The yeast protein secretion assays showed that transformants with SP-SUC2 or full-length Gas2 (Gas2^FL^-SUC2), but not Gas2 lacking the SP (Gas2^ΔSP^-SUC2), were able to grow on YPRAA medium and showed secreted invertase activity (Extended Data Fig.2d), indicating that the SP of Gas2 is functional in yeast. We also confirmed that Gas2 could be also translocated into plant cells during infection using a transformant of Δ*gas2* ectopically expressing Gas2-mCherry with a C-terminal nuclear localization signal (pRP27::Gas2-mCherry-NLS) (Extended Data Fig.2e). Furthermore, using an *M. oryzae* strain (Δ*gas2*/*GAS2-3HA*) expressing *GAS2-3HA* under the *EF1*α gene promoter^26^, we detected the Gas2-3HA protein both in the culture filtrate and mycelium (Fig. 1f), indicating that Gas2 could be secreted into culture medium. Finally, we generated transgenic rice plants expressing *3HA-GAS2* (*GAS2*-*OE*) in the Geng cultivar Nipponbare (NPB) (Extended Data Fig. 2f). Immunoblotting analysis showed that 3HA-Gas2 was distributed in both the cytoplasm and nuclei of transgenic rice cells (Fig. 1g). When considered together, these results indicate that Gas2 is a virulence protein that can be secreted into host plant cells and translocated into the nuclei of host cells during infection.

Since the *M. oryzae* Δ*gas2* mutant was attenuated in invasive hyphal growth^23^, we reasoned that Gas2 may play an important role in suppressing host immune responses. To test this, we used 3,3′-diaminobenzidine (DAB) staining to detect reactive oxygen species (ROS) in rice leaf sheath cells following infection with rice blast fungus. Notably, rice cells at 64.7% infection sites of the Δ*gas2* mutant were stained by DAB, a significantly higher percentage compared to 20.3% and 22.1% of the P131 and Δ*gas2*/*GAS2* strains, respectively (Extended Data Fig. 2g). RT-qPCR assays indicated that the plant defense-related genes *PBZ1* and *PR1* were more expressed in leaf sheaths infected by Δ*gas2* than by P131 and Δ*gas2*/*GAS2* strains (Extended Data Fig. 2h), suggesting that Gas2 may function as an effector to suppress plant immunity.

To verify that Gas2 is an effector shared by different types of rice fungal pathogens to suppress rice immunity, we compared symptoms on NPB and the *GAS2*-*OE* rice lines inoculated with different rice fungal pathogens. The *GAS2*-*OE* lines exhibited more severe symptoms than NPB after infection by WT strains of *M. oryzae* (Fig. 1h), *B. oryzae* (Fig. 1i), *U. virens* (Fig. 1j), *R. solani* (Fig. 1k) and *F. graminearum* (Fig. 1l), respectively. Moreover, the virulence of Δ*gas2* mutants of *M. oryzae* (Fig. 1h) and *B. oryzae* (Fig. 1i) was restored to the level of the isogenic WT strain on the *GAS2*-*OE* lines. Notably, the *GAS2*-*OE* lines showed significantly reduced induction of ROS and pathogenesis-related genes after chitin treatment, compared with NPB (Extended Data Fig. 2i,j). These results showed that the expression of Gas2 *in planta* suppresses PAMP-triggered immunity (PTI) in rice plants.

### Gas2 interacts with SnRK1β1A in the nuclei of plant cells

To examine how Gas2 suppresses PTI in rice, we conducted yeast two-hybrid screening (Y2H) of a cDNA library of *M. oryzae*-infected rice leaves using Gas2 as the bait. This screen identified several rice proteins that potentially interact with Gas2, including LOC_Os05g41220, which is a putative β1 subunit of the SNF1-related kinase 1 complex (SnRK1), hereafter referred to as SnRK1β1A. Similar to the Arabidopsis AKINβ1^27^, SnRK1β1A comprises a central carbohydrate-binding module (CBM) and a C-terminal conserved domain (βCTD) responsible for interacting with the α subunit of SnRK1 or other regulatory proteins (Extended Data Fig. 3a). Through Y2H assays, we revealed that Gas2 and SnRK1β1A interact with each other via the βCTD domain of SnRK1β1A and the DUF3129 domain of Gas2 (Fig. 2a, Extended Data Fig. 3b,c). Additionally, recombinant 2HA-Gas2 protein pulled down GST-SnRK1β1A protein, further supporting their physical interaction (Fig. 2b). In rice protoplasts, SnRK1β1A-cLuc co-immunoprecipitated with Gas2-3Flag, but not with GUS-3Flag (Fig. 2c). Luciferase complementation imaging (LCI) assays showed strong luciferase activity in *N. benthamiana* leaves co-expressing Gas2-nLuc and SnRK1β1A-cLuc (Extended Data Fig. 3d), supporting their interaction *in vivo*. Moreover, a bimolecular fluorescence complementation (BiFC) assay detected strong fluorescence signals in the nucleus but weak signals in the cytoplasm of *N. benthamiana* cells co-expressing Gas2 and SnRK1β1A (Fig. 2d), indicating that Gas2 interacts with SnRK1β1A in the nuclei of plant cells. Additionally, Y2H and LCI assays demonstrated that RsGas2, FfGas2, UvGas2, BoGas2, and FgGas2 interact with SnRK1β1A, whereas, BoGas2L and NcMAS1 do not (Fig. 2a, Extended Data Fig. 3e,f). Based on these results, we concluded that the Gas2 effectors from multiple rice fungal pathogens interact with SnRK1β1A.

**Fig. 2.**
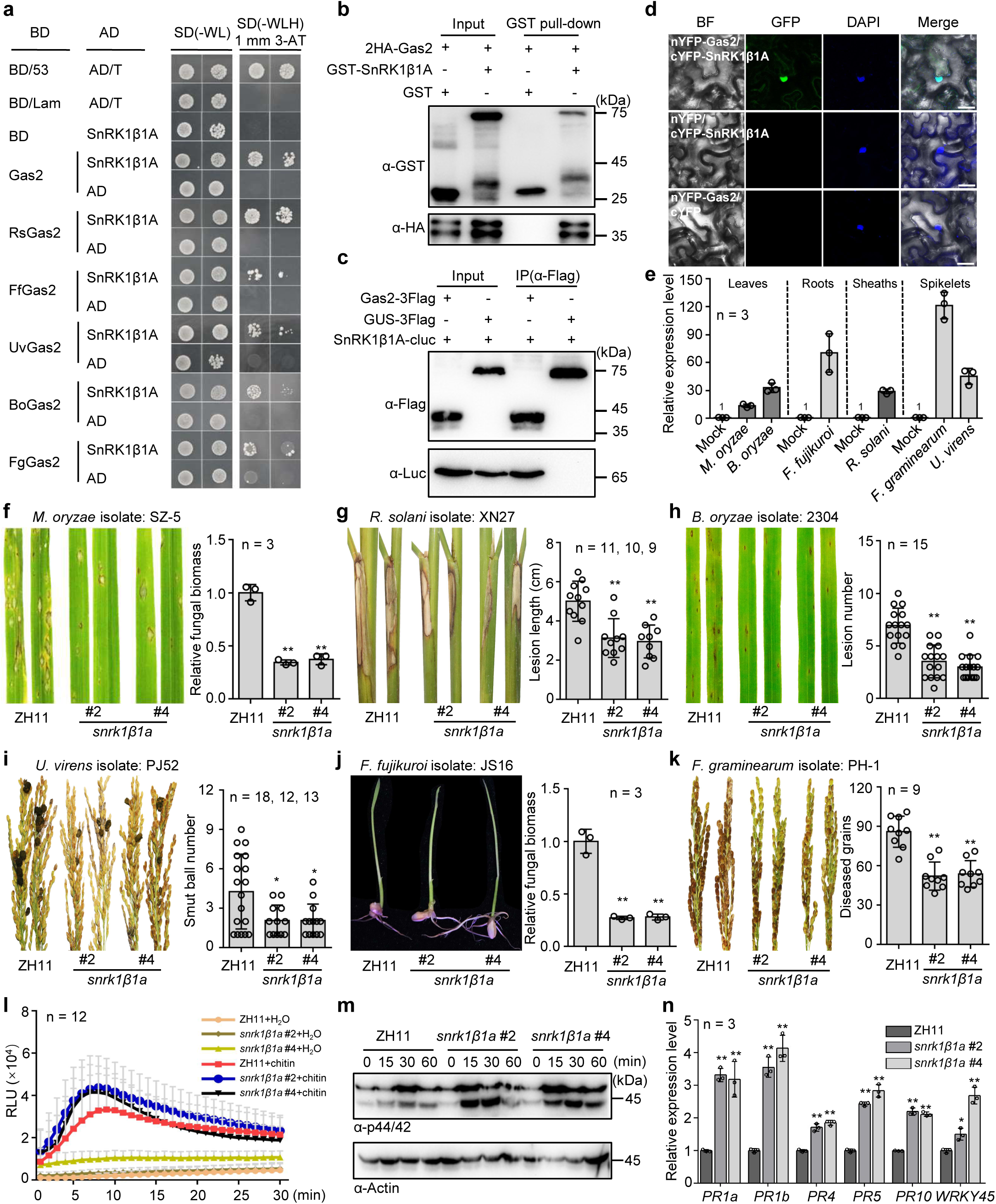
SnRK1β1A is targeted by Gas2 and negatively regulates rice immunity to multiple fungal pathogens. **a,** Yeast two-hybrid (Y2H) assays showed interaction of SnRK1β1A with Gas2 from *M. oryzae*, *R. solani*, *F. fujikuroi, U. virens, B. oryzae*, and *F. graminearum*. **b-d,** Interaction between SnRK1β1A and Gas2 from *M. oryzae* detected by GST pull-down (**b**), co-immunoprecipitation (co-IP) (**c**), and bimolecular fluorescence complementation (BiFC) (**d**). GST alone in (**b**) and GUS-3Flag in (**c**) served as negative controls. Scale bars in (**d**), 25 μm. **e,** *SnRK1*β*1A* was induced after infection by *M. oryzae*, *F. graminearum*, *F. fujikuroi*, *B. oryzae*, *R. solani* and *U. virens*. H_2_O treatment served as a control. **f-k,** Rice lines with disrupted *SnRK1*β*1A* (*snrk1*β*1a*) showed enhanced resistance as compared to its WT ZH11 to *M. oryzae* (**f**), *R. solani* (**g**), *B. oryzae* (**h**), *U. virens* (**i**), *F. graminearum* (**j**) and *F. fujikuroi* (**k**). Bar-graphs right to photos show disease severity or fungal biomass. **l,** Stronger ROS was induced in *snrk1*β*1a* lines upon chitin treatment. Rice leaf discs were immersed in 0.5 μM chitin for detection of ROS with the Luminol assay. **m, n,** *snrk1*β*1a* lines showed stronger activation of MAPK and induction of *PR* genes upon chitin treatment as compared with their WT ZH11, respectively. One-week-old rice seedlings were treated with 0.5 μM chitin for 0-60 minutes and then collected for phosphorylated MAPK detection with α-phospho-p44/42 antibody with α-actin as a loading control, for 6 hours to measure the expression of *PR* genes by RT-qPCR. Data represent Mean ± SD. Two-tailed Student’s t-test were used in **f-k**, **n**. **p < 0.01.

### SnRK1β1A negatively regulates rice immunity and its gene knockout confers broad-spectrum disease resistance

To assess the biological significance of the interaction of Gas2 with SnRK1β1A, we measured the expression of *SnRK1*β*1A* in different organs of rice, both with and without fungal infection. RT-qPCR analyses revealed that *SnRK1*β*1A* was expressed at low levels in the leaves, spikelets, roots and sheaths of healthy rice plants but significantly upregulated upon infection by diverse fungal pathogens (Fig. 2e, Extended Data Fig. 4a-f), suggesting that SnRK1β1A may play a crucial role in fungal invasion and colonization of plant tissue.

To explore the role of *SnRK1*β*1A*, we generated two independent knockout mutants in ZH11, a Geng rice cultivar, using the CRISPR-Cas9-mediated gene editing technique (Extended Data Fig. 4g). Infection assays showed that both the *snrk1*β*1a* mutants exhibited enhanced resistance to all tested *M. oryzae* strains (Fig. 2f, Extended Data Fig. 4h). The mutants were also resistant to *R. solani* (Fig. 2g), *B. oryzae* (Fig. 2h), *U. virens* (Fig. 2i), *F. fujikuroi* (Fig. 2j), and *F. graminearum* (Fig. 2k). Moreover, we detected significantly high-er levels of ROS (Fig. 2l), stronger MAPK activation (Fig. 2m) and up-regulated transcript levels of multiple PTI-related genes (Fig. 2n) in the *snrk1*β*1a* leaves following chitin treatment. Collectively, these findings indicate that SnRK1β1A acts as a negative regulator of rice immunity, and its gene knockout can confer broad-spectrum resistance to fungal diseases.

We finally conducted field trials at the Shangzhuang Experiment Station of CAU/Beijing, Panjin/Liaoning, Donggang/Liaoning and Enshi/Hubei to evaluate the effect of *SnRK1*β*1A* on rice disease resistance and growth under natural infection using mark-er-free *snrk1*β*1a* mutants (Extended Data Fig. 4i). As compared to ZH11, the mutant lines formed fewer and smaller blast lesions in leaves with significantly reduced disease indices in all test fields (Extended Data Fig. 4j). In addition, the *snrk1*β*1a* mutants also showed significantly reduced panicle blast, sheath blight and false smut in the Donggang field (Extended Data Fig. 4k-m). Importantly, the *snrk1*β*1a* lines displayed similar agronomic traits to ZH11, with the exception of heading, which occurred five days earlier in the mutants (Extended Data Fig. 5a-g). Overall, these results indicate that disrupting *SnRK1*β*1A* can confer broad-spectrum disease resistance without negatively impacting rice growth and yields.

### Gas2 protects SnRK1β1A from degradation

Since *GAS2*-*OE* lines exhibit enhanced susceptibility to multiple rice fungal diseases and SnRK1β1A negatively regulates rice immunity, we hypothesized that Gas2 may affect the accumulation of SnRK1β1A. To test this hypothesis, we first measured the SnRK1β1A-GFP protein levels in *N. benthamiana* leaves. As shown in Fig. 3a,b, much stronger SnRK1β1A-GFP fluorescence signals and hybridization signals were detected in tobacco cells co-expressing Gas2-3Flag compared to the cells expressing GUS-3Flag, despite similar transcript levels of *SnRK1*β*1A-GFP* in both cell types (Fig. 3b). These results suggest that Gas2 may enhance SnRK1β1A-GFP synthesis or prevent its degradation. Subsequently, we examined the SnRK1β1A-3Flag protein levels in rice protoplasts treated with cycloheximide (CHX), a protein synthesis inhibitor. In protoplasts co-expressing GFP with CHX, SnRK1β1A-3Flag levels were reduced, whereas SnRK1β1A-3Flag remained largely intact in protoplasts co-expressing Gas2-3HA (Extended Data Fig. 6a), indicating that Gas2 protects SnRK1β1A from degradation. Similarly, higher levels of SnRK1β1A-3Flag were detected in *GAS2-OE* rice protoplasts than in the WT rice protoplasts (Figure 3c). Furthermore, a cell-free degradation assay using recombinant GST-SnRK1β1A confirmed that over 50% of GST-SnRK1β1A proteins were degraded within 90 minutes when incubated with total protein extracts from ZH11 leaves and ATP. However, more than 60% of GST-SnRK1β1A proteins remained intact when recombinant 2HA-Gas2 protein was added (Extended Data Fig. 6b).

**Fig. 3.**
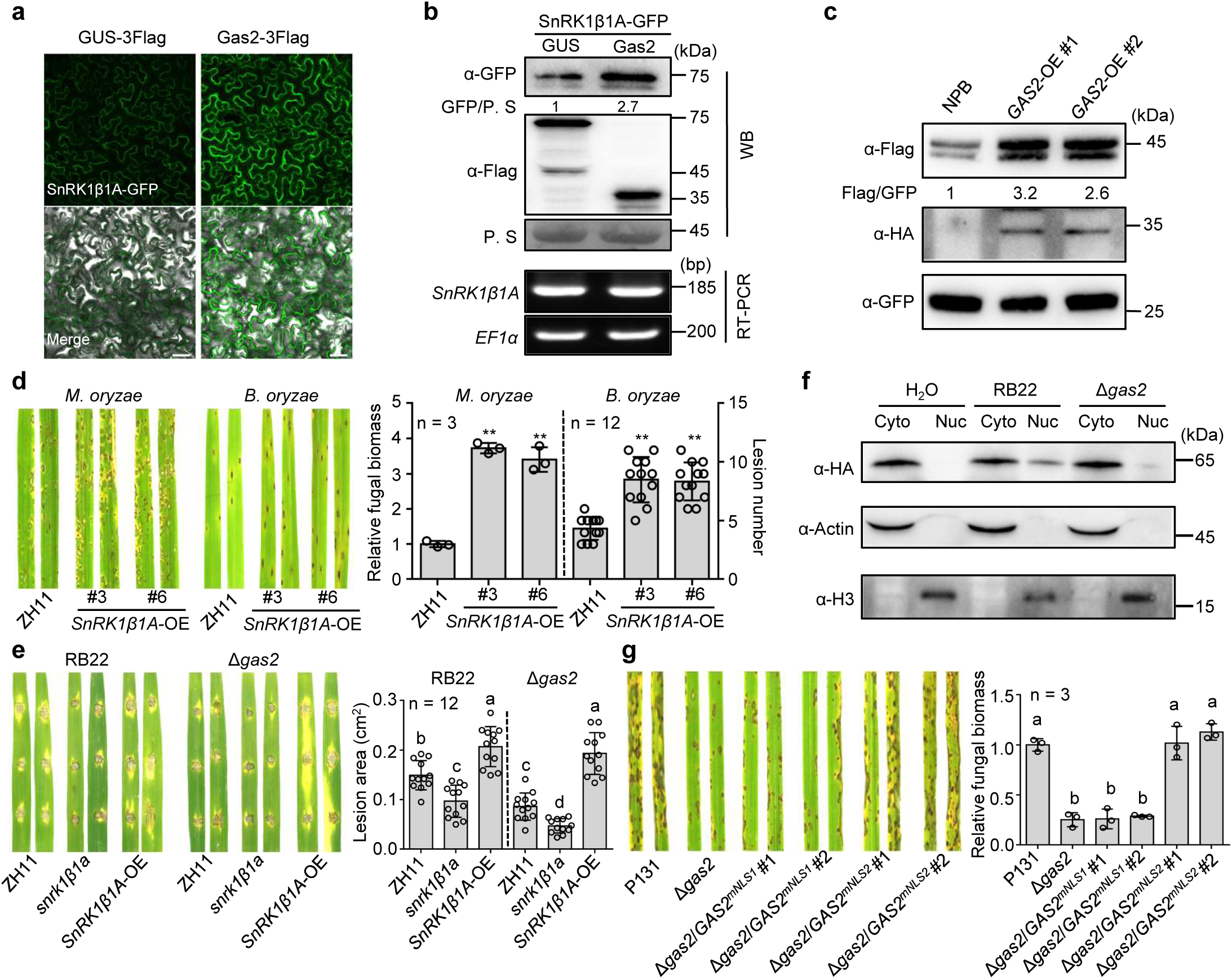
Gas2 inhibits degradation of SnRK1β1A and elicits SnRK1β1A nuclear localization. **a**,**b**, Higher levels of SnRK1β1A-GFP were detected in *N. benthamiana* leaves when transiently co-expressed with Gas2-3Flag but not withGUS-3Flag. Scale bars in (a), 25 μm. α-GFP and α-Flag antibodies were used for WB, with Ponceau S staining (P. S) as a loading control (Top). *SnRK1*β*1A* transcripts were detected using RT-PCR, with *NbEF1*α as an internal control (Bottom). **c,** SnRK1β1A-3Flag was more accumulated in *GAS2-OE* lines. Total proteins extracted from rice protoplasts co-expressing GFP were subjected to WB, which was conducted using α-Flag, α-HA and α-GFP antibodies. **d,** *SnRK1*β*1A-OE* lines were more susceptible to *M. oryzae* and *B. oryzae*. At 5 dpi, rice blast fungal biomass in was measured by qPCR, and brown spot lesion number per diseased leaf in was counted. **e,** Overexpression of *SnRK1*β*1A* in rice complements the defects of Δ*gas2*. The ZH11, *snrk1*β*1a* and *SnRK1*β*1A-OE* lines were drop-inoculated with the Δ*gas2* mutans and its WT strain RB22. Lesion areas were measured at 5 dpi using Image J software. **f,** Nuclear distribution of 3HA-SnRK1β1A in *SnRK1*β*1A*-OE lines was induced by WT strain but neither by the Δ*gas2* mutant of *M. oryzae* nor by H_2_O. α-HA antibody was used for the detection with α-Actin and α-H3 as the cytoplasmic and nuclear markers, respectively. **g,** Gas2 has two predicted nuclear localization signals (NLS), the first of which is essential for virulence. Gas2^mNLS1^ and Gas2^mNLS2^ are mutants with all arginine residues substituted by alanine in the predicted NLS motifs. Δ*gas2*/*GAS2^mNLS1^* and Δ*gas2*/*GAS2^mNLS2^* are transformants Δ*gas2* expressing Gas2^mNLS1^ and Gas2^mNLS2^, respectively. Fungal biomass was measured by qPCR at 5 dpi. The band intensities in (**b**,**c**) were quantified by Image J software. One-way ANOVA followed by post-hoc Turkey tests were used in (**e**,**g**), letters indicate significantly different groups; Two-tailed Student’s t-test was used in d. **p < 0.01.

To elucidate how Gas2 protects SnRK1β1A from degradation, we compared SnRK1β1A degradation in rice protoplasts under treatments with the 26S proteasome inhibitor MG132 and the autophagy inhibitor 3-Methyladenine, respectively. As shown in Extended Data Fig. 6c, MG132 but not 3-Methyladenine inhibited SnRK1β1A-3HA degradation, indicating that SnRK1β1A is likely degraded via the 26S proteasome system. Furthermore, we observed significantly higher ubiquitination of SnRK1β1A-3HA in tobacco cells co-expressing GUS-3Flag than those co-expressing Gas2-3Flag (Extended Data Fig. 6d). Since SnRK1β1A contains three lysine residues in its CTD domain that mediates its interaction with Gas2 (Extended Data Fig. 3b), we tested whether these residues serve as ubiquitination sites through point mutation. SnRK1β1A^K267R^, but neither SnRK1β1A^K248R^ nor SnRK1β1A^K275R^, exhibited reduced ubiquitination of SnRK1β1A, suggesting that K267 is a key ubiquitination site (Extended Data Fig. 6e). These findings collectively provide evidence that Gas2 may promote susceptibility by shielding SnRK1β1A from ubiquitination-mediated degradation.

Moreover, we generated transgenic rice lines overexpressing 3HA-SnRK1β1A in ZH11 (Extended Data Fig. 6f). Infection assays revealed that these transgenic rice lines developed more disease lesions with increased fungal biomass after infection by *M. oryzae* and *B. oryzae*, as compared to the WT ZH11 (Fig. 3d), confirming that *SnRK1*β*1A* is a susceptibility gene in rice. We then generated two Δ*gas2* mutants of *M. oryzae* strain RB22, virulent to ZH11 (Extended Data Fig. 6g). Infection assays showed that the Δ*gas2* mutants formed smaller lesions on both ZH11 and *snrk1*β*1a* leaves, compared to the isogenic strain RB22 while on the *SnRK1*β*1A-OE* leaves, the Δ*gas2* mutants formed lesions similar in size to those elaborated by RB22 (Fig. 3e), further supporting that SnRK1β1A is targeted by Gas2 to facilitate fungal infection.

### Gas2 can guide SnRK1β1A into the nucleus of plant cell

Given that SnRK1β1A and Gas2 predominantly interact in the nucleus of plant cells (Figure 2d), and that SnRK1β1A is distributed in both the cytoplasm and nucleus in the absence of Gas2 (Figure 3a), we hypothesized that Gas2 may promote the nuclear localization of SnRK1β1A. To investigate this potential protein translocation, we quantified the proportion of *N. benthamiana* cells displaying nuclear SnRK1β1A-GFP signals. As shown in Extended Data Fig. 7a, approximately 50% of tobacco cells co-expressing Gas2-3Flag exhibited nuclear signals of SnRK1β1A-GFP, compared to only 5% in the control tobacco cells co-expressing GUS-3Flag. Western blot analysis further confirmed nuclear localization of SnRK1β1A-GFP in tobacco cells co-expressing Gas2-3Flag, but not in those co-expressed co-expressing GUS-3Flag (Extended Data Fig. 7b). We also examined whether the subcellular localization of SnRK1β1A is influenced by rice blast fungus infection. In leaves of *SnRK1*β*1A*-overexpressing (OE) plants inoculated with RB22, the 3HA-SnRK1β1A protein was detected in both nuclear and cytoplasmic extracts. However, in mock-treated leaves or those inoculated with the Δ*gas2* mutant, SnRK1β1A was found exclusively in the cytoplasm (Figure 3f). These results collectively suggest that Gas2 is crucial for directing SnRK1β1A to the nucleus.

Since Gas2 is detectable in the nucleus of rice cells (Figure 1g), we speculated it might contain a nuclear localization signal (NLS). By using the APBS routine in PyMol (http://www.pymol.org/), we identified two motifs enriched with basic amino acids in Gas2, designated as NLS1 and NLS2 (Extended Data Fig. 7c). To determine the role of these two NLSs in Gas2 localization, we generated Gas2^mNLS1^ and Gas2^mNLS2^ mutants by substituting all lysine/arginine residues with alanine (Extended Data Fig. 7d) and observed subcellular localization of the Gas2 mutants in *N. benthamiana* cells. As shown in Extended Data Fig. 7e, Gas2^mNLS2^ was distributed in both the nucleus and cytoplasm, while Gas2^mNLS1^ was predominantly localized in the cytoplasm, suggesting that NLS1 functions as a nuclear localizing signal. Since NLS1 resides in the DUF3129 domain of Gas2, which mediates the interaction with SnRK1β1A, we further tested whether NLS1 affects this interaction. As shown in Extended Data Fig. 7f,g, Gas2^mNLS1^ failed to interact with SnRK1β1A, indicating that the basic amino acids in NLS1 are critical not only for the nuclear localization of Gas2 but also for its interaction with SnRK1β1A. Moreover, we introduced Gas2^mNLS1^ and Gas2^mNLS2^ vectors into the Δ*gas2* strain to assess their function. Infection assays showed that Δ*gas2*/*GAS2^mNLS2^*restored WT virulence, whereas Δ*gas2*/*GAS2^mNLS1^* failed to rescue the WT phenotype (Fig. 3g). These results indicate that NLS1 is essential for the virulence function of Gas2.

### SnRK1β1A interacts and functions antagonistically with SnRK1**α**1

To elucidate how SnRK1β1A facilitates rice susceptibility, we examined its interaction with SnRK1α1, the catalytic subunit of SnRK1, which is known to positively regulate PTI in rice^2,28^. The Y2H and LCI assays demonstrated that SnRK1α1 interacts with SnRK1β1A in yeast cells (Extended Data Fig. 8a) and tobacco leaves (Extended Data Fig. 8b), respectively. Furthermore, *in vitro* pull-down assays confirmed that recombinant 2Flag-SnRK1α1 protein binds explicitly to GST-SnRK1β1A (Extended Data Fig. 8c). Additionally, a BiFC assay verified the interaction between SnRK1β1A and SnRK1α1, with fluorescence signals predominantly localized in the cytoplasm of *N. benthamiana* cells (Extended Data Fig. 8d). Collectively, these results indicate that SnRK1β1A interacts with SnRK1α1 in the cytoplasm.

To further explore how SnRK1β1A regulates the function of SnRK1α1 through this interaction, we generated two independent *snrk1*β*1asnrk1*α*1* double mutants, named *snrk1*β*1asnrk1*α*1* #6 and #10 (Extended Data Fig. 8e), in the background of previously reported *snrk1*α*1 #1* mutant^29^. As expected^2,28^, the *snrk1*α*1* mutants exhibited increased susceptibility to *M. oryzae* (Extended Data Fig. 8f). Notably, the *snrk1*β*1asnrk1*α*1* double mutants, exhibited similar susceptibility of the *snrk1*α*1* mutant to *M. oryzae* (Fig. 4a). Moreover, the double mutants displayed reduced chitin-induced ROS bursts, similar to the levels detected in the *snrk1*α*1* lines (Fig. 4b). Thus, the loss of *SnRK1*α*1* abolishes the enhanced resistance conferred by *snrk1*β*1a*, suggesting that SnRK1β1A functions antagonistically with SnRK1α1 in regulating rice immunity.

**Fig. 4.**
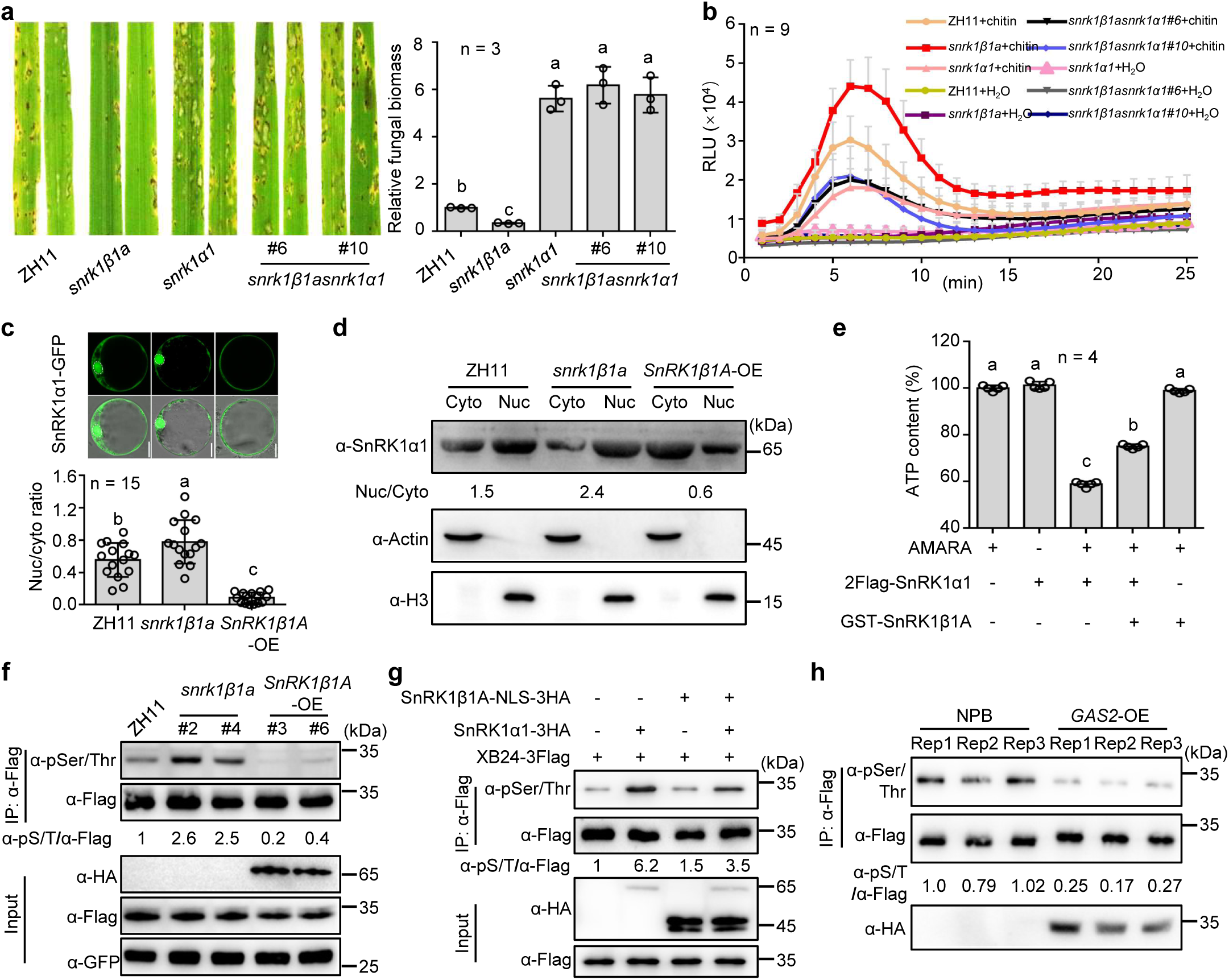
SnRK1β1A negatively regulates SnRK1α1 functions. **a,** Disruption of *SnRK1*α*1* abolished *snrk1*β*1a*-mediated blast resistance. ZH11 and its mutants were inoculated with *M. oryzae* isolate SZ-5. Fungal biomass was measured by qPCR at 5 dpi. **b,** The *snrk1*β*1asnrk1*α*1* mutant showed less ROS accumulation than *snrk1*β*1a* upon chitin treatment. **c,** More nuclear localizations of SnRK1α1-GFP transiently expressed in rice protoplasts of *snrk1*β*1a* than in those of ZH11, and *SnRK1*β*1A*-OE. Scale bars, 10 μm. Bar-graph showing the nuclear to cytoplasmic fluorescence ratios of SnRK1α1-GFP. **d,** The cytoplasmic and nuclear distributions of endogenous SnRK1α1 in ZH11, *snrk1*β*1a* and *SnRK1*β*1A*-*OE* lines. α-SnRK1α1 antibody was used for the detection, with α-Actin and α-H3 as the cytoplasmic and nuclear markers, respectively. Nuclear to cytoplasmic fluorescence ratios in **c** and **d** were estimated by Image J software. **e,** SnRK1β1A inhibited the kinase activity of SnRK1α1. Both the proteins were expressed and purified from *Escherichia coli*. AMARA peptide was used as a substrate. **f,** SnRK1β1A inhibited SnRK1α1-mediated XB24 phosphorylation. *XB24-3Flag* plasmid is expressed in protoplasts of ZH11, *snrk1*β*1a* and *SnRK1*β*1A*-OE lines. 3HA-SnRK1β1A in *SnRK1*β*1A*-OE lines was detected using α-HA antibody. GFP was used as a co-expression control. **g,** XB24-3Flag phosphorylation was reduced in rice protoplasts co-expressing SnRK1α1-3HA and SnRK1Aβ1A-NLS-3HA as compared those expressing SnRK1α1-3HA. **h,** Reduced XB24-3Flag phosphorylation in the rice protoplasts of *GAS2*-OE line as compared to its WT NPB. 3HA-Gas2 in *GAS2*-OE lines was detected using α-HA antibody. XB24-3Flag in (**f**-**h**) was immunoprecipitated for detecting its phosphorylation with α-pSer/Thr antibody. The band intensities in (**f**-**h**) were quantified by Image J software. One-way ANOVA followed by post-hoc Dunnett tests were used in (**a**) or by Turkey tests were used in (**c**,**e**); letters indicate significantly different groups.

### SnRK1β1A restricts SnRK1**α**1 kinase activity

Previous studies have reported that SnRK1α1s primarily function in the nucleus in Arabidopsis and barley^30,31^. However, our observations reveal that rice SnRK1β1A interacts with SnRK1α1 in the cytoplasm (Extended Data Fig. 7d). This led us to hypothesize that the cytoplasmic interaction between SnRK1β1A and SnRK1α1 may limit the nuclear localization of SnRK1α1. To test this hypothesis, we examined the subcellular localization of SnRK1α1-GFP in epidermal cells of transiently expressed *N. benthamiana* leaves. In control cells co-expressing RFP, SnRK1α1-GFP signals were observed in both the nucleus and cytoplasm. However, in cells co-expressing SnRK1β1A-RFP, GFP signals were predominantly localized in the cytoplasm (Extended Data Fig. 8g). This observation was confirmed by nuclear and cytoplasmic protein fractionation from tobacco leaves expressing SnRK1α1-3Flag with or without SnRK1β1A-3Flag (Extended Data Fig. 8h). Furthermore, in rice protoplast transfection assays, SnRK1α1-GFP signals were detected in both the nucleus and cytoplasm in ZH11 protoplasts, with a greater nuclear proportion observed in *snrk1*β*1a* protoplasts, but less frequently in the nucleus of *SnRK1*β*1A*-OE protoplasts (Fig. 4c). Western blot analyses corroborated these findings, showing that SnRK1α1 predominantly accumulates in the nucleus of *snrk1*β*1a* plants and in the cytoplasm of *SnRK1*β*1A-OE* lines (Fig. 4d). Collectively, these data provide evidence that SnRK1β1A inhibits the nuclear localization of SnRK1α1.

To test whether SnRK1β1A can directly affect the kinase activity of SnRK1α1, we performed an *in vitro* assay using the AMARA peptide, a well-established AMPK substrate in mammalian cells^32^. As shown in Fig. 4e, recombinant 2Flag-SnRK1α1 protein phosphorylates the AMARA peptide and consumes ATP, while 2HA-SnRK1β1A alone does exhibit the kinase activity. However, the kinase activity of 2Flag-SnRK1α1 was significantly reduced in the presence of 2HA-SnRK1β1A. Further, we assessed the *in vivo* impact of SnRK1β1A on SnRK1α1 by assessing the phosphorylation level of XB24, a rice ATPase targeted and phosphorylated by SnRK1α1 using a rice protoplasts system^28^. A much higher phosphorylation level of XB24-3Flag was detected in *snrk1*β*1a* lines than in ZH11, where- as the phosphorylation of XB24-3Flag was barely detectable from *SnRK1*β*1A-OE* lines (Figure 4f). Taken together, SnRK1β1A impedes the kinase activity of SnRK1α1 *in vitro* and *in vivo*.

Y2H and LCI assays indicated that Gas2 is unable to directly interact with SnRK1α1 (Extended Data Fig. 9a,b) while BiFC assays showed that the interaction of SnRK1β1A and SnRK1α1generated stronger fluorescence signals in the nucleus of *N. benthamiana* cells co-expressing Gas2-3Flag (Extended Data Fig. 9c). To assess the impact of nuclear translocation of SnRK1β1A, we gauged the phosphorylation level of XB24 in rice protoplasts^28^. XB24-3Flag exhibited a high phosphorylation level in rice protoplasts co-expressing *SnRK1*α*1-3HA* but a significantly lower phosphorylation level in rice protoplasts co-expressing *SnRK1*α*1-3HA*/*SnRK1*β*1A-NLS-3HA*, which harbors a C-terminal NLS (Fig. 4g), indicating that SnRK1β1A also inhibits the activity of SnRK1α1 in the nucleus. These results collectively suggest that SnRK1β1A is normally localized in the cytoplasm and that during *M. oryzae* infection, Gas2 is secreted into rice cells, guiding the translocation of SnRK1β1A from the cytoplasm into the nucleus, where SnRK1β1A inhibit SnRK1α1 activity. Consistently, phosphorylation levels of XB24 in the *GAS2*-*OE* line were significantly lower than that in WT rice (Figure 4h), indicating that Gas2 in rice cells can result in lower SnRK1 kinase activity.

## Discussion

Many *S* genes have been identified across diverse plant species through mutant screening or as targets of pathogen-specific effectors^6^. While disrupting these genes often results in resistance to a specific pathogen^7,12,21^, achieving broad-spectrum disease resistance to multiple pathogen classes through *S* gene disruption remains arduous due to associated penalties in growth, developmental, and/or yield^16,20^. Although some *S* genes can confer wide-spectrum disease resistance without negative effects^17,18,33^, an effective strategy to identify such *S* genes remains elusive. Here, we identified SnRK1β1A in rice as an S protein by leveraging Gas2, an effector conserved among multiple rice fungal pathogens, including *M. oryzae*, *R. solani*, *U. virens*, *F. fujikuroi*, *F. graminearum*, and *B. oryzae*. We also showed that disrupting *SnRK1*β*1A* in rice enhanced resistance to the diseases caused by these fungal pathogens. Notably, *SnRK1*β*1A* is expressed at low levels in healthy rice plants but is highly induced upon infection by these pathogens. Its disruption activates PTI responses only upon infection, without incurring growth and yield penalties. Together, our study demonstrates that broad-spectrum disease resistance can be achieved without compromising plant fitness by inactivating infection-inducible *S* genes that are convergently targeted by effectors conserved in multiple pathogens.

SnRK1 is a conserved plant serine/threonine kinase complex, analogous to SNF1 in yeast and AMPK in animals^34^, comprising α1, β1 and γ1 subunits and functioning as a cell energy sensor in response to environmental stresses and nutritional limitations^35^. While the roles of the α1 subunits in plants have been extensively studied, revealing their involvement in nutrient sensing, stress responses, metabolic balance, and immunity against various pathogens^36^, the functions of the β1 subunits have received less attention. In this study, we found that rice *SnRK1*β*1A* mutants displayed no obvious defect in growth, development, and yield, mirroring the findings from *A. thaliana* mutants of the β1 subunit gene *AKIN*β*1*^37^. Remarkably, our results showed that the *snrk1*β*1a* lines became resistant to a broad range of fungal pathogens, while disrupting *SnRK1*α*1* in a *snrk1*β*1a* mutant abolished this disease resistance, indicating that SnRK1β1A functions by inhibiting SnRK1α1. Our investigation further revealed that SnRK1β1A inhibits the kinase activity of SnRK1α1 on substrates, such as XB24, a protein crucial for the PTI response to false smut caused by *U. virens*^29^. In addition, the disease-resistant *snrk1*β*1a* lines showed a higher degree of nuclear localization of SnRK1α1, while the disease-susceptible *SnRK1*β*1A*-OE lines exhibited reduced nuclear localization of the protein. This underscores the critical role of SnRK1β1A in regulating the nuclear localization of SnRK1α1 to confer disease resistance. Together, our findings provide multiple lines of evidence that SnRK1β1A negatively regulates the catalytic subunit SnRK1α1 in rice, whose overexpression confers broad-spectrum disease resistance with growth and yield penalties^2^. However, further studies are required to verify whether SnRK1β1A orthologues in other plant species and other β subunits of SnRK1 in rice also act as negative regulators of plant immunity and SnRK1α1 activity. Moreover, we revealed that SnRK1β1A accumulates and undergoes reduced ubiquitination in plant cells expressing the *M. oryzae* effector Gas2, suggesting that ubiquitination-mediated degradation may be an important regulatory mechanism for SnRK1β1 function. Further investigation is required to assess whether SnRK1β1 subunits are similarly regulated by ubiquitination in other eukaryotes, including yeast and animals.

MAS proteins, characterized by a conserved DUF3129 domain, are widely distributed in fungi. Although first identified in *Blumeria graminis* f. sp. *hordei*^24^, MAS proteins have been reported as virulence factors in several other fungal pathogens, including *M. oryzae*^23,38^, *Verticillium dahliae*^39^, *Sclerotinia sclerotiorum*^40^ and *Podosphaera xanthii*^25^. However, not all MAS proteins in pathogenic fungi are required for virulence^38^, and some non-pathogenic fungi also harbor MAS proteins, suggesting that virulence-associated MAS proteins possess distinct features yet to be fully characterized. Here, we revealed that Gas2, a MAS protein conserved across multiple rice fungal pathogens, convergently targets rice SnRK1β1A to promote susceptibility. Furthermore, we identified a positively charged patch within the DUF3129 domain of Gas2, which functions as a nuclear localization signal and is required for its interaction with SnRK1β1A. Additionally, we demonstrated that the α helix at the extreme C-terminus of Gas2 is essential for its virulence function, although this region is not required for its interaction with SnRK1β1A. These features appear to be characteristic of Gas2-type MAS proteins. However, further research is needed to identify key motifs or amino acids in Gas2 that mediate its interaction with SnRK1β1A, and to explore the features of other virulence-associated MAS proteins.

## Methods

### Plant materials and growth conditions

The rice cultivar Zhonghua 11 (ZH11, *japonica*) was used as wild type (WT) to generate *SnRK1*β*1A* knockout and *SnRK1*β*1A* overexpression in this study. The previously reported *snrk1*α*1* mutant^29^ was used to generate *snrk1*β*1asnrk1*α*1* double mutant. The rice cultivars Nipponbare (NPB, *japonica*) served as the WT for *GAS2* heterogeneous transformation and rice fungal pathogenicity assays in this study. The rice cultivar Lijiangxintuanheigu (LTH, *japonica*) was used for *M. oryzae* pathogenicity assays. All rice and transgenic plants were grown in a greenhouse at 28°C with 70% relative humidity under a 12-hour light/12-hour dark cycle. Rice seedlings were grown on sterilized tubes with solid half-strength Murashige and Skoog (MS) medium in a growth chamber at 28°C for protoplast preparation.

Barley (*Hordeum vulgare* cv E9) and tobacco (*Nicotiana benthamiana*) plants were grown at 25°C under long-day conditions (16 hour day/8 hour night) in a greenhouse for *M. oryzae* infection assays and transient expression assays, respectively.

### Fungal pathogens

The rice blast fungus *M. oryzae* field isolates P131, RB22, SZ-5, SZ-4 and SZ-2^41^ were cultured on oat tomato agar (OTA) plates at 28°C. The floral pathogen *U. virens* isolates JS60-2 and PJ52, and the brown spot fungus 2304 were grown on potato sucrose (PS) plates and sporulated on RSEDOMA plates (20 g·L^-1^ rice straw, 20 g·L^-1^ dextrose, 20 g·L^-1^ oatmeal and 20 g·L^-1^ agar) at 28°C. The sheath blight fungus *R. solani* isolate XN27, the head blight fungus *F. graminearum* isolates PH-1, and the bakanae pathogen *F. fujikuroi* isolate JS16 were all maintained on potato dextrose agar (PDA) plates at 28°C in this study.

### Construction of transgenic rice plants

The coding sequence (CDS) of *GAS2* (without the signal peptide) and *SnRK1*β*1A* fused with 3×HA at the N-terminal were inserted into the vector pCAMBIA1301, which contains the maize *UBIQUTIN1* promoter and the hygromycin B phosphotransferase gene (Hyg resistance). The resulting constructs pCAMBIA1301-*3HA-GAS2* and pCAMBIA1301-*3HA-SnRK1*β*1A* were transformed into NPB and ZH11 plants, respectively. Positive transgenic seedlings were selected with hygromycin (100 mg·L^-1^), and genomic DNA was extracted for PCR detection of the hygromycin B phosphotransferase gene.

To generate CRISPR/Cas9 mediated gene knockout mutants, 20-bp gene-specific guide RNA sequences targeting *SnRK1*β*1A* were synthesized, annealed, and ligated into pOs-sgRNA, which was subcloned into the vector pBGK023. The constructs were transformed into rice plants through *Agrobacterium tumefaciens*-mediated transformation. The mutation sites of all the target genes were verified by PCR-based sequencing.

### Fungal gene knockout and transformation

A homologous substitution approach was used to generate the *GAS2* mutant in *M. oryzae* isolates P131 and RB22, *U. virens* isolate JS60-2, *B. oryzae* isolate 2304, *F. graminearum* isolate PH-1 and *F. fujikuroi* isolate JS16. Approximately 1.5 kb flanking sequences of *GAS2*, *UvGAS2*, *BoGAS2*, *FgGAS2* or *FfGAS2* were constructed as homologous arms onto knockout vector pKOV21. Subsequently, protoplasts of individual WT strains were isolated and transformed by the PEG/CaCl_2_ method as previously described^26^. All transformants were screened by PCR to detect flanking regions of the hygromycin marker using specific primers, and Δ*gas2* mutants were further confirmed by Southern blot analysis. Probes used for Southern blot were labeled with the DIG High Prime DNA Labeling and Detection Starter Kit II.

For Δ*gas2* complementation assays, the CDS of *GAS2*, *GAS2* mutants and *GAS2*-orthologs from multiple fungi with promoter fragments of *GAS2* were amplified and cloned into the vector pGTN, which contains the neomycin resistance gene. For Gas2 secretion assays, the CDS of *GAS2* with the *EF1*α promoter^26^ was amplified and cloned into the vector pKN-3HA. For Gas2 translocation assays, the CDS of *GAS2* fused with *mCherry-NLS* was amplified and cloned into the vector pRKN with an *RP27* promotor. All resulting vectors were transformed into protoplasts of the Δ*gas2* mutant. The transformants were screened by PCR and assayed for phenotype complementation. For Δ*Fggas2* complementation assays, the CDS of *GAS2* orthologs with promoter fragments of *GAS2* were amplified and cloned into the vector pKN-GFP and then transformed into Δ*Fggas2* protoplasts.

### Fungal pathogen inoculation on rice plants

For *M. oryzae* inoculation assays, all isolates were cultured on OTA plates at 28°C to produce spores. Rice leaves were sprayed with the conidial suspensions at a concentration of 5 × 10^4^ spores·mL^-1^ in 0.025% Tween 20. The inoculated leaves were incubated in a moist, dark chamber at 28°C for 24 h and then grown under regular illumination. Disease lesions were photographed at 5 days post-inoculation (dpi). The relative fungal biomass was measured with DNA-based quantitative PCR (qPCR) using the cycle threshold (CT) value of the *M. oryzae MoPot2* gene against the CT of the rice *ubiquitin* gene.

For the *M. oryzae* infection process assessment, the indicated strains were injected into the hollow interior of rice sheaths and barley epidermis. Microscopy observations were performed using sheaths infected with P131, Δ*gas2* and Δ*gas2*/*GAS2* at indicated times under a Nikon 90i microscope. The sheaths infected with Δ*gas2*/*GAS2* strain and barley epidermis infected with Δ*gas2*/*GAS2-mCherry-NLS* strain were observed at 30 hours post-inoculation with a fluorescence microscope. DAPI staining indicated cell nuclei.

For rice false smut inoculation assays, *U. virens* isolates JS60-2 and PJ52 were grown on PS plates for 2 weeks and then cultured in PS medium at 28°C with shaking at 140 rpm for 7 days. The mixture of hyphae and spores were blended and diluted to 1 × 10^6^ spores·mL^-1^ in PS medium as the inoculum. The inoculum of JS60-2 and PJ52 were injected into rice spikes of NPB and ZH11 background, respectively, with a syringe at 5-7 days before rice heading. False smut balls formed on rice panicles were counted 30 dpi.

For sheath blight inoculation assays, *R. solani* isolate XN27 was cultured on PDA plate for 2 d at 28°C. Square filter paper (5 mm × 5 mm) was sterilized and co-incubated with fungal for 3 days in a new plate. The fungi on filter paper were then inserted into the fourth leaf sheath at the tillering stages. The length of sheath blight lesions was measured at 10 dpi.

For brown spot inoculation assays, *B. oryzae* isolates were cultured on PDA plate for 3 days at 28°C and then grown on RSEDOMA plates for 7 days at 28°C under cool white fluorescent lights for conidiation. Rice leaves were sprayed with the conidial suspensions at a concentration of 1 × 10^4^ spores·mL^-1^ in 0.025% Tween 20. The inoculated leaves were incubated in a moist, dark chamber at 28°C for 24 h and then grown under regular illumination. Disease lesions were photographed, and the number of lesions was counted at 5 dpi.

For head blight inoculation assays, discs of all *F. graminearum* strains mycelia growing on PDA plate were cultured in carboxymethylcellulose sodium (CMC) medium at 28°C with shaking at 150 rpm for 2 days. The mixture was filtered and diluted to 2 × 10^5^ spores·mL^-1^ in sterile H_2_O as the inoculum. PH-1 was injected into rice spikes for resistance evaluation of *snrk1*β*1a* mutants; The inoculum of PH-1 and Δ*Fggas2* was injected into ZH11 rice spikes for virulence test. The diseased grains were photographed and counted at 7 dpi. The relative fungal biomass was measured with DNA-based qPCR using the CT of the *F. graminearum FgCHS5* gene against the CT of the rice *ubiquitin* gene.

For rice bakanae fungus inoculation assays, discs of *F. fujikuroi* JS16 mycelia growing on PDA plates were cultured in CMC medium at 28°C with shaking at 150 rpm for 2 days. The mixture was filtered and diluted to 1 × 10^6^ spores·mL^-1^ in sterile H_2_O, which was co-incubated with rice seeds that had been pregerminated for 24 h. Inoculated seeds were continued to germinate for 5 days. The relative fungal biomass was measured by DNA-based qPCR using the CT of the *F. fujikuroi 28S* ribosomal DNA gene against the CT of the rice *ubiquitin* gene.

All inoculation assays were repeated independently at least three times.

### Yeast secretion assay

The yeast secretion assay was performed as previously described ^42^. The CDS encoding putative signal peptide (SP) of Gas2 (22-aa SP), the CDS encoding Gas2 without SP (Gas2^ΔSP^) and the CDS encoding full-length Gas2 (Gas2^FL^) were fused in frame with the CDS encoding SUC2 lacking the SP. Avr1b and Mg87 served as the positive and negative controls, respectively. The constructs were then transformed into the yeast-invertase-deficient strain YTK12 using the PEG/LiAc transformation method. The transformants were cultured on CMD-W medium and YPRAA medium. The invertase enzymatic activity was detected by the reduction of 2, 3, 5-triphenyltetrazolium chloride (TTC) to insoluble 1, 3, 5-triphenylformazan (TPF) with red coloration. Transformants were cultured in liquid CMD/-W medium and then collected and resuspended in buffer (10 mM acetic acid–sodium acetate buffer (pH 4.7):10 % sucrose solution (w/v): sterile distilled water=2:1:3) at 37°C for 10 minutes. Then, the supernatant, after centrifugation, was put into a glass test tube containing 0.1% TTC solution for 5 minutes. The picture was immediately obtained after the colorimetric change.

### Yeast two-hybrid (Y2H) assay

Y2H screening was carried out to identify Gas2 interacting proteins using a rice cDNA library. The CDS of *GAS2* without SP was cloned into pGBKT7 as the bait vector. For Y2H assay, the matchmaker GAL4 two-hybrid system (Clontech) was used. The CDS of WT *GAS2*, *GAS2* NLS mutants, *GAS2* homologues and *SnRK1*α*1* were cloned in plasmid pGBKT7 as the bait vectors, and the CDS of *SnRK1*β*1A*, and the truncated *SnRK1*β*1A* were cloned in plasmid pGADT7 as the prey vectors. The bait plasmids and the corresponding prey plasmids were co-transformed into the yeast Y2H Gold strain, following the manufacturer’s instructions. Positive transformants were selected to grow on SD/-Leu-Trp-His medium with different concentrations of 3-Amino-1,2,4-Triazole (3AT).

### Luciferase complementation imaging (LCI) assay

The LCI assay was conducted as previously described^43^. Briefly, the CDS of WT *GAS2*, *GAS2 NLS* mutants, *GAS2* homologues and *SnRK1*β*1A* were cloned into plasmid pCAMBIA1300-HA-nLUC, and the CDS of *SnRK1*β*1A*, *SnRK1*α and the truncated *SnRK1*β*1A* were cloned in plasmid pCAMBIA1300-cLUC.The resulting plasmids were transformed into the *A. tumefaciens* strain GV3101 using the freeze-thaw method. Two corresponding vectors were co-agroinfiltrated into 4-week-old *N. benthamiana* leaves with an induction buffer (10 mM MES, pH 5.6, 10 mM MgCl_2_, and 100 mM acetosyringone). Luciferase activity was assessed at 48 hours post-agroinfiltration by spraying 1 mM of D-luciferin onto the infiltrated *N. benthamiana* leaves. The luminescence signals were captured by a CCD imaging apparatus. Next, leaf discs (4 mm in diameter) were punched from infiltrated leaf areas and placed in a 96-well plate containing 100 μL of sterile water. After replacing the sterile water with 100 μL of 1 mM D-luciferin, leaf discs were kept in the dark for 5 minutes. Luminescence was then quantified using a microplate reader with an integration time of 0.5 s. Finally, proteins were extracted from infiltrated leaves by extraction buffer (25 mM Tris-HCl, pH 7.5, 150 mM NaCl, 1 mM EDTA, 10% glycerol, 1% Triton X-100, 0.5% NP40) and detected by western blot analyses using anti-HA and anti-Luc antibodies to confirm the expression of indicated proteins. All primers are listed in Table 1.

### Co-immunoprecipitation (co-IP) assay

Ten-day-old etiolated rice seedlings grown on 1/2 MS medium in the dark were used for protoplast isolation. The *35S*-*SnRK1*β*1A*-*cluc* plasmid was co-transfected with *35S*-*GAS2*-*3Flag* or *35S*-*GUS*-*3Flag* plasmid into rice protoplasts via PEG-mediated transformation. The transfected protoplasts were incubated for 16 h in the dark and then collected for protein extraction. The extracted proteins were incubated with anti-FLAG M2 affinity beads in IP buffer (50 mM Tris-HCl, pH 7.5, 150 mM NaCl, 1 mM MgCl_2_, 1% Triton X-100) at 4°C for 4 h on a rotating shaker. The beads were then washed 3–5 times with pre-cooled 1×PBS buffer, and the immunoprecipitates were eluted using 1×SDS–PAGE loading buffer. The samples were subjected to immunoblot analyses with anti-Flag and anti-Luc antibodies.

### Bimolecular fluorescence complementation (BiFC) assay

The CDS of *GAS2* and *SnRK1*β*1A* were cloned into plasmid pUC19 with a C-terminal nYFP, and the CDS of *SnRK1*β*1A* and *SnRK1*α*1* were cloned into plasmid pUC19 with a C-terminal cYFP as previously described^44^. Agrobacterium strains transformed with each construct or empty vector were mixed at OD_600_ of 0.5 and then infiltrated into 4-week-old *N. benthamiana* leaves. Green fluorescence was observed in *N. benthamiana* leaves at 36 hours post infiltration using laser scanning confocal microscopy (Leica SP8). DAPI staining indicates nuclei.

### GST pull-down assay

The CDS of *GAS2* fused with *2×HA* tag at the N-terminal and *SnRK1*α*1* fused with *2×Flag* tag at the N-terminus were cloned into plasmid pETM-13. The CDS of *SnRK1*β*1A* was cloned into plasmid pGEX with an N-terminal GST tag. The expression and extraction of GST-SnRK1β1A, 2×HA-Gas2-6×His, and 2×Flag-SnRK1α1-6×His proteins from *E. coli* were performed as previously described^45^. Briefly, recombinant protein extracts were stored in binding buffer (20 mM Tris, 150 mM NaCl, 5 mM DTT, 4 mM EDTA, pH 7.4, and 1% Triton X-100). Anti-GST beads (30 μL) were washed three times in the binding buffer. One milliliter of the binding buffer and 10 μg proteins were separately added to anti-GST beads and incubated at 4°C for 3 h with constant rotation. The pellets were washed five times with the binding buffer and boiled for 10 minutes for immunoblotting analyses with anti-GST, anti-HA or anti-Flag antibodies.

### RT-PCR and RT-qPCR

To quantify the expression levels of *GAS2*, *SnRK1*β*1A* or *PR* genes upon treatment with rice fungal pathogens or chitin, samples (rice leaves, roots, sheaths or spikelets) were collected at indicated times. Total RNA was extracted using the KK Fast Plant RNA Kit (ZOMANBIO). Then 1 μg of RNA was reverse-transcribed to cDNA with the 1st Strand cDNA Synthesis Kit (Vazyme) according to the manufacturer’s instructions. RT-PCR assays were performed with 2×Taq HiFi PCR mix (Mei5Bio) to detect the expression of *GAS2* and *SnRK1*β*1A* in rice and tobacco, respectively. RT-qPCR assays were performed with 2×SYBR Green Pro Taq HS Premix (Accurate). The rice *Actin* gene (*LOC_Os03g50885*) and the *Nicotiana benthamiana EF1*α gene served as the internal control.

### Chitin-induced ROS burst, MAPK activation and PR gene expression

For ROS burst assay, the leaves of three-week-old rice seedlings were punched with 4 mm diameter puncher and kept in water overnight before the assays. Leaf discs were soaked in 100 μL solutions with luminol substrate, peroxidase and chitin (0.5 μM), or sterile ddH_2_O as control. Luminescence was recorded at 1-minute intervals for 45 minutes with a Spectra Max i3x luminometer.

For the MAPK activation assay, one-week-old cultured rice seedlings were kept in water overnight and then treated with chitin (0.5 μM) or sterile H_2_O plus 0.01% Silwet L-77. The seedlings were collected at 0, 15, 30 and 60 minutes after treatment. The total protein was extracted with the extraction buffer (150 mM NaCl, 25 mM Tris-HCl pH 7.5, 1 mM EDTA, 10% glycerin, 0.5% NP-40, 1% Triton X-100, 10mM DTT, 1% PMSF and 0.1% protease inhibitor cocktail) including phosphatase inhibitor PhosSTOP (Roche), and then detected by immunoblotting with the phospho-p44/42 MAPK and anti-Actin antibodies.

For the quantification of *PR* gene expression, cultured rice seedlings treated with chitin (0.5 μM) were collected at 6 hours post-treatment. Total RNA was extracted from those samples, and the PR gene expression in those samples was quantified using RT-qPCR.

### Protein degradation assay *in planta* and cell-free system

For the SnRK1β1A degradation assay in rice protoplasts, the plasmid *SnRK1*β*1A-3Flag* was either transfected alone or co-transfected with the *GAS2-3HA* construct into rice protoplasts prepared from leaf sheaths of 10-day-old ZH11 seedlings. An empty vector, 35S-GFP, was co-expressed in each reaction as an internal reference. After 14 hours of incubation in the dark, the protoplasts were treated with 50 μM of the protein synthesis inhibitor cycloheximide (CHX). The abundance of SnRK1β1A protein was detected at 0 and 4 hours post-CHX treatment using anti-HA, anti-Flag and anti-GFP antibodies. The band intensities were quantified using Image J software.

For the SnRK1β1A degradation assay in tobacco leaves, the plasmid *SnRK1*β*1A-3HA* was transiently expressed in the tobacco leaves for 36 hours. Total proteins were extracted at 4 hours post-CHX treatment, along with treatment with the proteasome inhibitor MG132 or the autophagy inhibitor 3-Methyladenine (3-MA), and were determined by immunoblotting analyses with anti-HA antibody. The band intensities were quantified using Image J software. Ponceau S staining (P. S) served as a loading control.

The cell-free SnRK1β1A degradation assay was conducted as described previously^42^. Briefly, in vitro purified GST-SnRK1β1A recombinant protein with or without 2HA-Gas2 recombinant protein was incubated with the crude protein extract isolated from ZH11 rice leaves at 25°C for 0, 45, and 90 minutes. A final concentration of 10 mM ATP was added to the mixture. The reaction was halted by boiling for 5 minutes in the SDS sample buffer. Western blots were performed to detect the remaining GST-SnRK1β1A proteins in these reaction mixtures using an anti-GST antibody. Anti-actin antibody served as a loading control.

### Nuclear-cytoplasmic fractionation assay

The nuclear-cytoplasmic fractionation assay was conducted according to previously established protocols with some modifications^46^. Briefly, plant tissues (*N. benthamiana* or rice leaf) weighing 1 g were collected at specified time points post-treatment, ground into fine powders, and then homogenized in 2 mL of extraction buffer I (20 mM Tris-HCl, pH 7.5, 20 mM KCl, 2.5 mM MgCl_2_, 2 mM EDTA, 25% glycerol, 250 mM sucrose, 1×Protease Inhibitor Cocktail, and 5 mM DTT). The resulting lysate was filtered through a triple layer of Miracloth to remove debris, and the filtrate was centrifuged at 1,500 rpm for 5 minutes to pellet nuclei. The supernatant was transferred into a new tube and centrifuged at 8,000 rpm for 10[minutes. The resulting supernatant was collected as the cytoplasmic fraction. The nuclei containing pellet were resuspended in 5 mL of extraction buffer II (20 mM Tris-HCl, pH 7.4, 25% glycerol, 2.5 mM MgCl_2_, and 0.2% Triton X-100), followed by centrifugation for 10 minutes at 1,500 rpm. This step was repeated for 5 cycles. The resulting pellets were then resuspended in 500 μL of extraction buffer III (20 mM Tris-HCl, pH 7.5, 0.25 M sucrose, 10 mM MgCl_2_, 0.5% Triton X-100, and 5 mM β-mercaptoethanol).

The nuclear fraction was carefully layered on the top of 500 μL of extraction buffer IV (20 mM Tris-HCl, pH 7.5, 1.7 M sucrose, 10 mM MgCl_2_, 0.5% Triton X-100, 1×Protease Inhibitor Cocktail, and 5 mM β-mercaptoethanol), and then centrifuged at 13,000 rpm for 45 minutes. The resulting pellet was resuspended in 200 μL extraction buffer I. All procedures were performed on ice or at 4°C. Samples were detected via western blot assays using anti-HA, anti-Flag and anti-SnRK1α1 antibodies. anti-Actin and anti-Histone H3 antibodies served as markers for the cytoplasmic and nuclear fractions, respectively. The band intensities were quantified using Image J software.

### *In vivo* ubiquitination assay

For the *in vivo* SnRK1β1A ubiquitination assay, SnRK1β1A-3HA was transiently expressed in *N. benthamiana* leaves either with Gas2-3Flag or GUS-3Flag. To identify ubiquitination site on SnRK1β1A, WT SnRK1β1A, and mutants SnRK1β1A^K248R^-3HA, SnRK1β1A^K267R^-3HA and SnRK1β1A^K275R^-3HA were transiently expressed in *N. benthamiana* leaves. Protein extracts were immunoprecipitated with an EZview^TM^ Red Anti-HA Affinity Gel, and the level of ubiquitination was detected by immunoblot analysis with an α-Ubiquitin antibody. Ponceau S staining (P. S) served as a loading control.

### SnRK1 kinase assays

The *In vitro* kinase assay was performed using purified 2Flag-SnRK1α1 and GST-SnRK1β1A proteins expressed in *E. coli*. The AMARA polypeptide served as a general substrate. Reactions were initiated by adding kinase buffer (50 mM Tris HCl, pH 7.5; 10 mM MgCl_2_, 0.1 mM EGTA, 1 mM DTT and 50 μM ATP), and then incubated for 30 minutes at 25°C. SnRK1α1 activity was measured using the Kinase-Lumi^TM^ Plus Luminescent Kinase Assay Kit (Beyotime). Briefly, reactions were added to 96 well plate and immediately co-incubated with equal Kinase-Lumi^TM^ solution for 10 minutes. Luminescence was quantified after using a microplate reader with an integration time of 0.25 s. The ATP contents in each reaction was calculated.

Additionally, *in vivo* SnRK1 kinase activity in the rice protoplasts was detected by XB24 phosphorylation, as previously described^28^. The plasmid XB24-3Flag was transfected or co-transfected with plasmids SnRK1α1-3Flag, SnRK1β1A-3Flag or SnRK1β1A-NLS-3Flag into rice protoplasts. Extracted proteins from the protoplasts were incubated with EZview^TM^ Red Anti-Flag M2 affinity Gel to immunoprecipitate XB24-3Flag. The phosphorylation of XB24-3Flag was detected by immunoblotting with anti-pSer/Thr antibody. The band intensities were quantified using Image J software. The *in vivo* SnRK1 kinase assays were repeated independently three times.

### Subcellular localization and quantitative analysis

The subcellular localization of Gas2, Gas2 mutants, SnRK1β1A and SnRK1α1 was analyzed in *Nicotiana benthamiana* and rice protoplasts. The CDS of *GAS2* and its mutated forms, *SnRK1*β*1A* and *SnRK1*α*1* were inserted into the pCAMBIA1300 plasmid with a C-terminal GFP, which contains the 35S promoter. The resulting constructs were agroinfiltrated into *N. benthamiana* leaves were transformed into rice protoplasts. Expression of fusion proteins was observed with a laser scanning confocal microscopy (Leica SP8). In addition, quantitative analyses of SnRK1α1-GFP nuclear/cytoplasmic ratio were determined on the confocal images using ImageJ software.

### Field trials of marker-free *snrk1*β*1a* lines

Field evaluation of rice blast and sheath blight disease resistance was performed at CAU-Shangzhuang Crop Experiment Station/Beijing, Panjin/Liaoning, Donggang/Liaoning and Enshi/Hubei. Among them, Enshi has been used as a famous disease nursery because of the high incidence rate of rice blast^16,20,47^. In brief, approximately 30 rice plants of each line were grown in the field, all of which are surrounded by highly susceptible rice. Rice leaf blast disease and rice false smut disease were investigated at tillering stage using a 9-level classification system (Chinese Agricultural Industry Standards, NY/T 3685-2020) and a 5-level classification system^48^, respectively, which is then calculated as a disease index. In addition, rice plants grown in a neighbor non-diseased paddy field in the same area were assayed to observe agronomic traits.

### Phylogenetic analysis

All sequences of Gas2 and Gas2-homologous proteins from multiple fungi or SnRK1β1 subunits from rice, barley and Arabidopsis were obtained from the NCBI database using pBLAST. Multiple sequence alignments of proteins were done in MEGA X and a phylogenetic tree of aligned sequences was also constructed in MEGA X using the Maximum Likelihood method^49^.

### Protein structure analysis

The structure models of Gas2 and Gas2-homologous proteins were predicted using AlphaFold2^50^. The analysis of protein structure superposition was displayed using the PyMOL software (https://pymol.org/2/).

### Primers

Primer sequences are provided in Supplementary Table 1-5.

### Quantification and statistical analysis

Quantification analysis of lesion length, fungal growth, kinase activity, gene expression, H_2_O_2_ accumulation, grain production and other measurements was conducted in GraphPad Prism 6. All values are presented with mean ± SD. The exact values of n are indicated in the legends. The significance of the difference was examined by two-tailed Student’s t test or One-way ANOVA followed by post-hoc Turkey tests.

### Reporting summary

Further information on research design is available in the Nature Research Reporting Summary linked to this paper.

### Data availability

All data are available within this Article and its Supplementary Information. Original gel blots are shown in Supplementary Information Fig. 1. Original data points in graphs and statistical analyses of this study are shown in the Source Data files. Source data are provided with this paper.

## Acknowledgment

This study was supported by the National Natural Science Foundation of China (Grant No. 32293244 to Y.-L.P.), the China Agricultural Research System (CARS-1-43 to J.Y.), and Pingduoduo-China Agricultural University joint project (Grant no. PC2023A01005 to Y.-L.P.), and 2115 Talents project from China Agricultural University (to Y.-L.P.). We are grateful to Shen Chen for providing *B. oryzae* 2304 isolate; Wenxian Sun for providing *U. virens* PJ52 and JS60-2 isolates; Dao-Xiu Zhou for providing *snrk1*α*1* mutants; Nicholas J. Talbot for critical review and constructive discussion; Jin-Rong Xu and Sheng-Yang He for their suggestions and discussions.

## Author contributions

Y.-L.P., G.Y., and X.L. conceived the project. G.Y., X.W., M.L., Z.H. and F.Z. performed the experiments. S.W., and X.Z. predicted and analyzed the structures of MAS proteins. Y.-L.P., G.Y., X.L., X.W., J.Y., H.G., V.B., W.Zhu, W.Zhao. analyze the data. Y.-L.P., G.Y., X.L., and V.B. wrote the paper. All authors discussed and commented on the manuscript.

## Extended Data Figure legends

**Extended Data Fig. 1.**
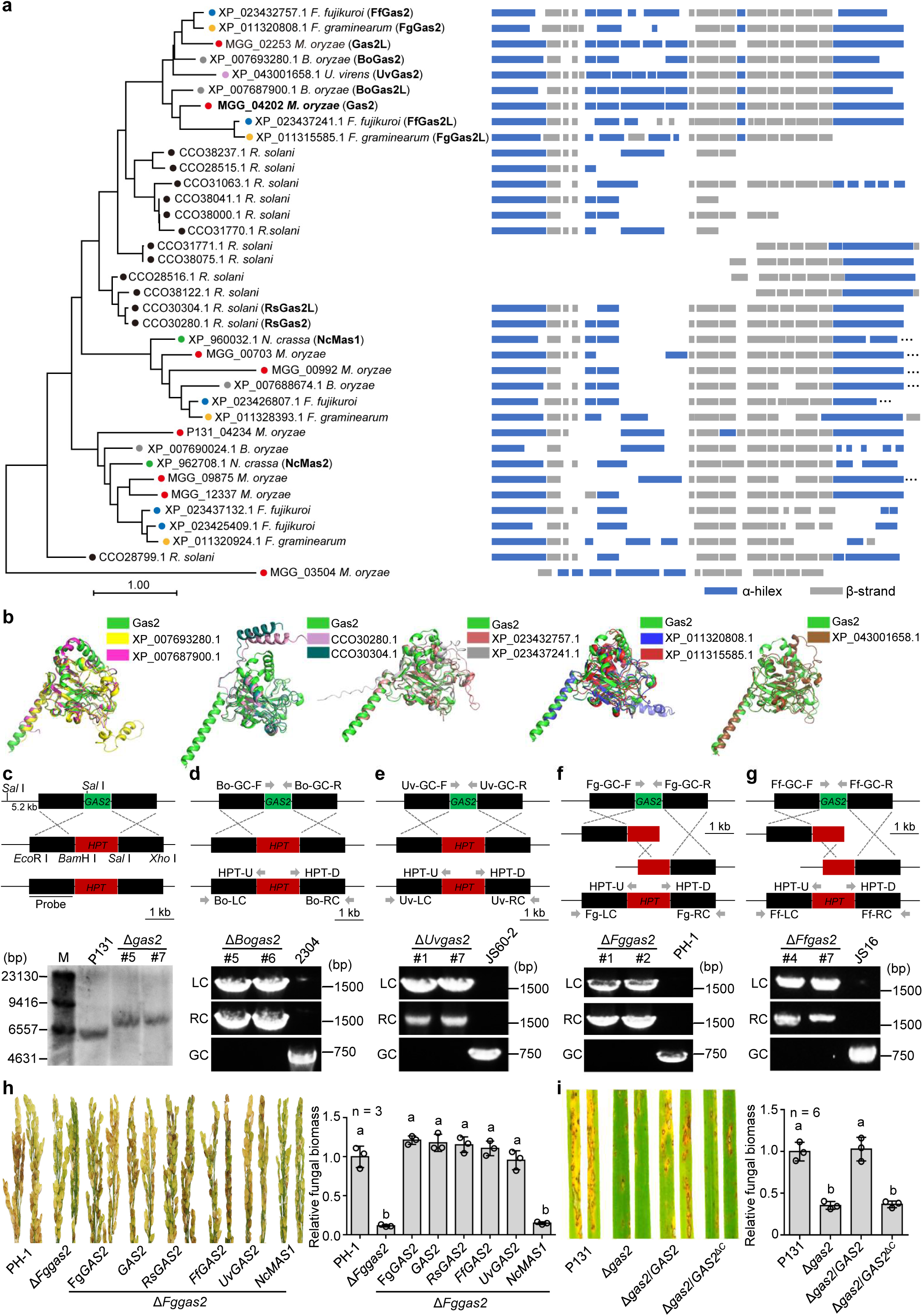
Gas2 is conserved in rice fungal pathogens. **a,** Phylogenetic analysis and secondary structures of Gas2-homologous proteins. The phylogenetic tree of Gas2 homologous proteins was constructed in MEGA X using the Maximum Likelihood method. The secondary structures of Gas2 homologous proteins were predicted with PSIPRED 4.0. **b,** Comparisons of the AlphaFold2 predicated 3D-structure of Gas2 from *M. oryzae* with its orthologues from *Bipolaris oryzae* (XP_007693280.1/BoGas2 and XP_007687900.1/BoGas2L), *Rhizoctonia solani* (CCO30280.1/RsGas2 and CCO30304.1/RsGas2L), *F. fujikuroi* (XP_023432757.1/FfGas2 and XP_023437241.1/FfGas2L), *Fusarium graminearum* (XP_011320808.1/FgGas2 and XP_011315585.1/FgGas2L) and *Ustilaginoidea virens* (XP_043001658.1/UvGas2), which cause brown spot, sheath blight, bakanae, head blight and false smut in rice, respectively. **c-g,** Schematic diagrams and verification of Δ*gas2* (c), Δ*Bogas2* (d), Δ*Uvgas2* (e), Δ*Fggas2* (f) and Δ*Ffgas2* (g) knockout mutants. Δ*gas2* was checked according to Southern blot analysis, genomic DNA isolated from wild-type (WT) P131 and the two Δ*gas2* mutants were digested with *SalI*. The blot analysis was performed by the digoxin kit instructions. *HindIII*-digested λ-DNA was used as a marker (c). Genomic DNA isolated from the other fungal mutants and its corresponding WT strains were amplified by PCR using specific primers (d-g). **h,** Virulence of *F. graminearum* Δ*Fggas2* mutant on Nipponbare (NPB) could be rescued by *GAS*2-orthologues but unrelated *MAS* from saprophytic *N. crassa*. Relative fungal biomass was measured by qPCR at 5 dpi. **i,** the WT virulence of Δ*gas2* mutant could not be rescued by introducing *GAS2* without the C-terminus (Δ*gas2*/*GAS2*^Δ*C*^). Rice leaves were sprayed with *M. oryzae* P131, Δ*gas2* and Δ*gas2*/*GAS2*^Δ*C*^ strains. Relative fungal biomass was measured by qPCR at 5 dpi. Data represent mean ± SD. One-way ANOVA followed by post-hoc Turkey tests were used in **h**,**i**, letters indicate significantly different groups.

**Extended Data Fig. 2.**
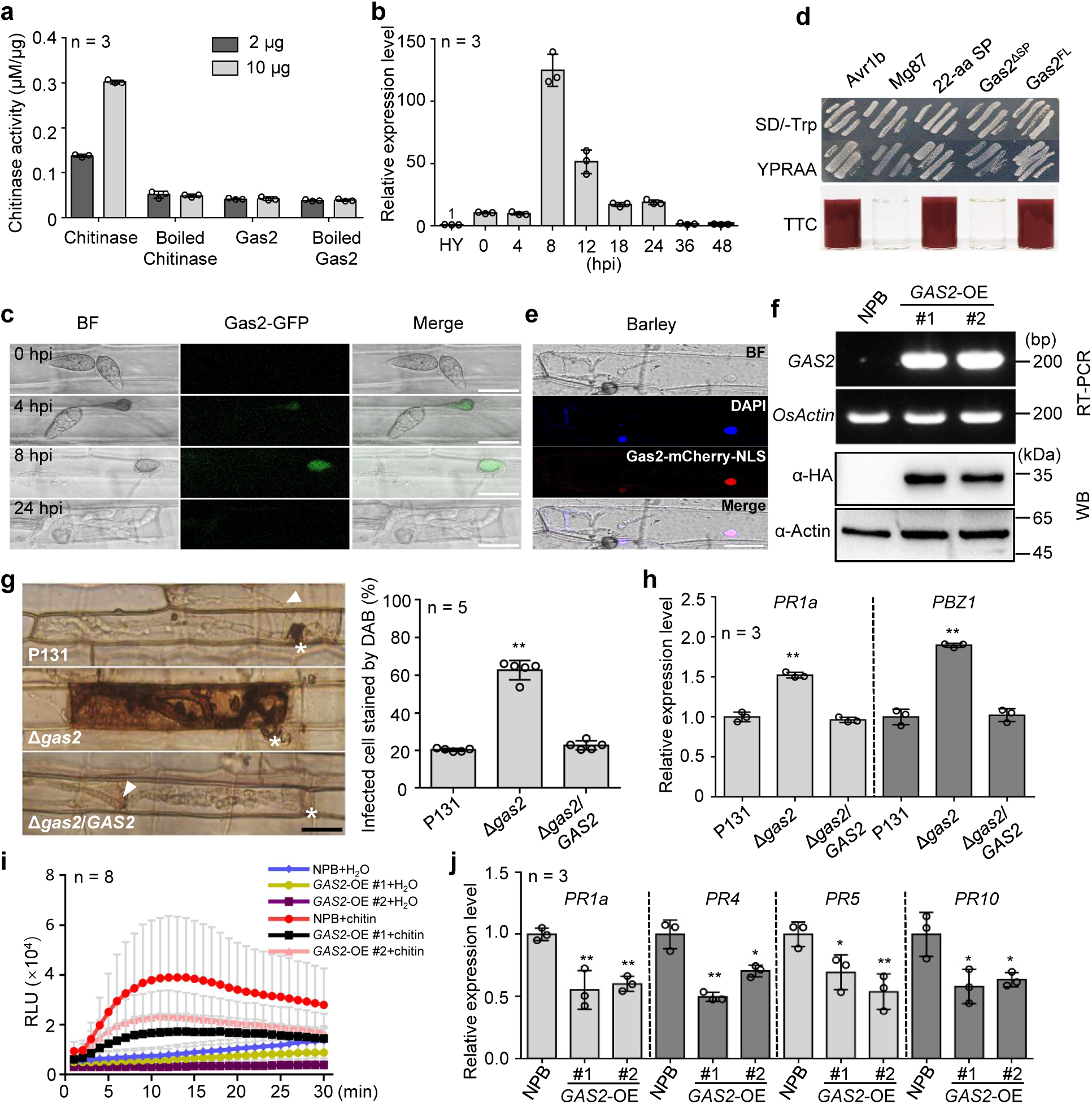
Gas2 is expressed during the appressorial penetration and suppresses PTI in rice plants. **a,** Gas2 lacks chitinase activity. 2HA-Gas2 purified in *E. coli* was used to detect chitinase activity. Chitinase from *Oryzae sativa* was used as a positive control and the boiled chitinase was used as a negative control. **b,** *GAS2* is specifically expressed during the appressorial penetration. Relative expression of *GAS2* normalized to *MoActin* was calculated using RT-qPCR with RNA extracted from hyphae and rice leaves inoculated with *M. oryzae* strain P131 at indicated times. **c,** Gas2 localized to the appressorium in rice leaf sheaths. Microscopic observation of Gas2-GFP driven by the native promoter in rice cells at indicated time points. Scale bar, 20 μm. **d,** The signal peptide from Gas2 could lead to the secretion of *SUC2* to allow growth of yeast transformants on SD-Trp medium and YPRAA medium (Top) and secretion of invertase into liquid medium (Bottom). **e,** Gas2-mCherry-NLS could be observed in the nucleus of barely epidermal cells infected by an *M. oryzae* transformant expressing Gas2-mCherry-NLS at 30 hours post-inoculation (hpi). Gas2-mCherry signals clearly overlapped with DAPI staining. Scale bar, 20 μm. **f,** Transgenic rice lines of NPB overexpressing *3HA-GAS2* (GAS2-OE) verified at transcriptional (top) and protein (bottom) levels. RNA and protein were extracted from indicated rice leaves for RT-PCR assays and western blot analysis with α-HA antibody, respectively. *Actin* was used as an internal control. **g,** Rice leaf sheath cells infected with Δ*gas2* showed stronger ROS accumulation as compared with the cells infected with P131 and complementary strain Δ*gas2*/*GAS2*. At 36 hpi, rice leaf sheath cells were stained with 3,3′-diaminobenzidine (DAB) and infected cells stained with DAB were counted. Asterisks indicate appressoria, and arrows represent invasive hyphae invading neighboring cells. Scale bars, 25 μm. Two-tailed Student’s t-test was used, **p < 0.01. **h,** Rice leaf sheath cells infected with Δ*gas2* showed stronger induction of *PR* genes as compared with the cells infected with P131 and complementary strain Δ*gas2*/*GAS2*. RNA samples were collected from rice leaf sheaths inoculated with indicated strains at 36 hpi. *OsActin* was used as an internal control. **i,** *GAS2*-OE lines showed weaker ROS burst upon chitin treatment as compared with its WT NPB. Rice leaf discs were immersed in 0.5 μM chitin for detection of ROS with the Luminol assay. **j,** *GAS2*-OE lines showed weaker induction of *PR* genes upon chitin treatment as compared with its WT NPB. One-week-old rice seedlings were treated with 0.5 μM chitin for 6 hours to measure the expression of *PR* genes by RT-qPCR. Two-tailed Student’s t-tests were used in (**g**,**h**,**j**), *p < 0.05, **p < 0.01.

**Extended Data Fig. 3.**
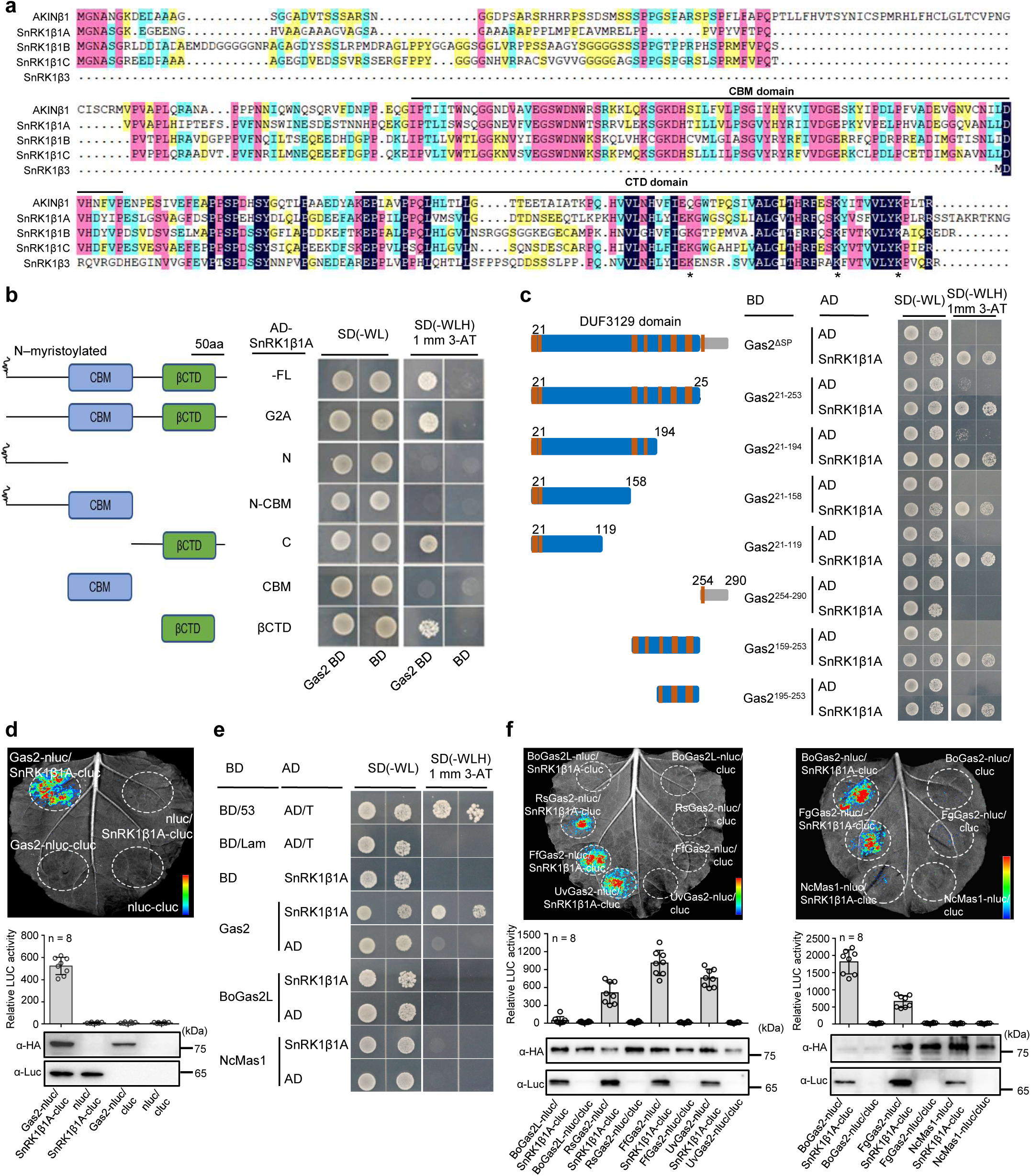
Gas2 from multiple rice fungal pathogens interact with SnRK1β1A. **a,** The alignment of Arabidopsis AKINβ1, rice SnRK1β1A, SnRK1β1B, SnRK1β1C and SnRK1β3. Lines indicate the CBM domain and βCTD domain, and asterisks indicate the conserved lysine residues in rice SnRK1 β subunits that may serve as potential ubiquitination sites. **b,** Y2H assays showed interaction of Gas2 with the CTD domain of SnRK1β1A. The curve in the N-terminal indicates potential myristoylated glycine (G), the blue and green boxes indicate the CBM and CTD domains, respectively. **c,** Y2H assays showed interaction of SnRK1β1A with DUF3129 domain of Gas2. The blue box and the orange box indicate the DUF3129 domain and the β-strand of Gas2, respectively. **d,** LCI assays showed the interaction of SnRK1β1A with Gas2 from *M. oryzae*. The relative luciferase activity was measured and protein expression was confirmed by immunoblot analysis with α-HA and α-Luc antibodies. **e,** Y2H assays showed interaction of SnRK1β1A with Gas2 from *M. oryzae*, but neither with Gas2-like protein from *B. oryzae* nor with unrelated MAS proteins from *N. crassa*. **f,** LCI assays showed interaction of SnRK1β1A with Gas2 from *B. oryzae*, *U. virens, R. solani*, *F. fujikuroi* and *F. graminearum*, but neither with Gas2-like protein from *B. oryzae* nor with unrelated MAS proteins from *N. crassa*. The relative luciferase activity was measured, and protein expression was confirmed by immunoblot analysis with α-HA and α-Luc antibodies.

**Extended Data Fig. 4.**
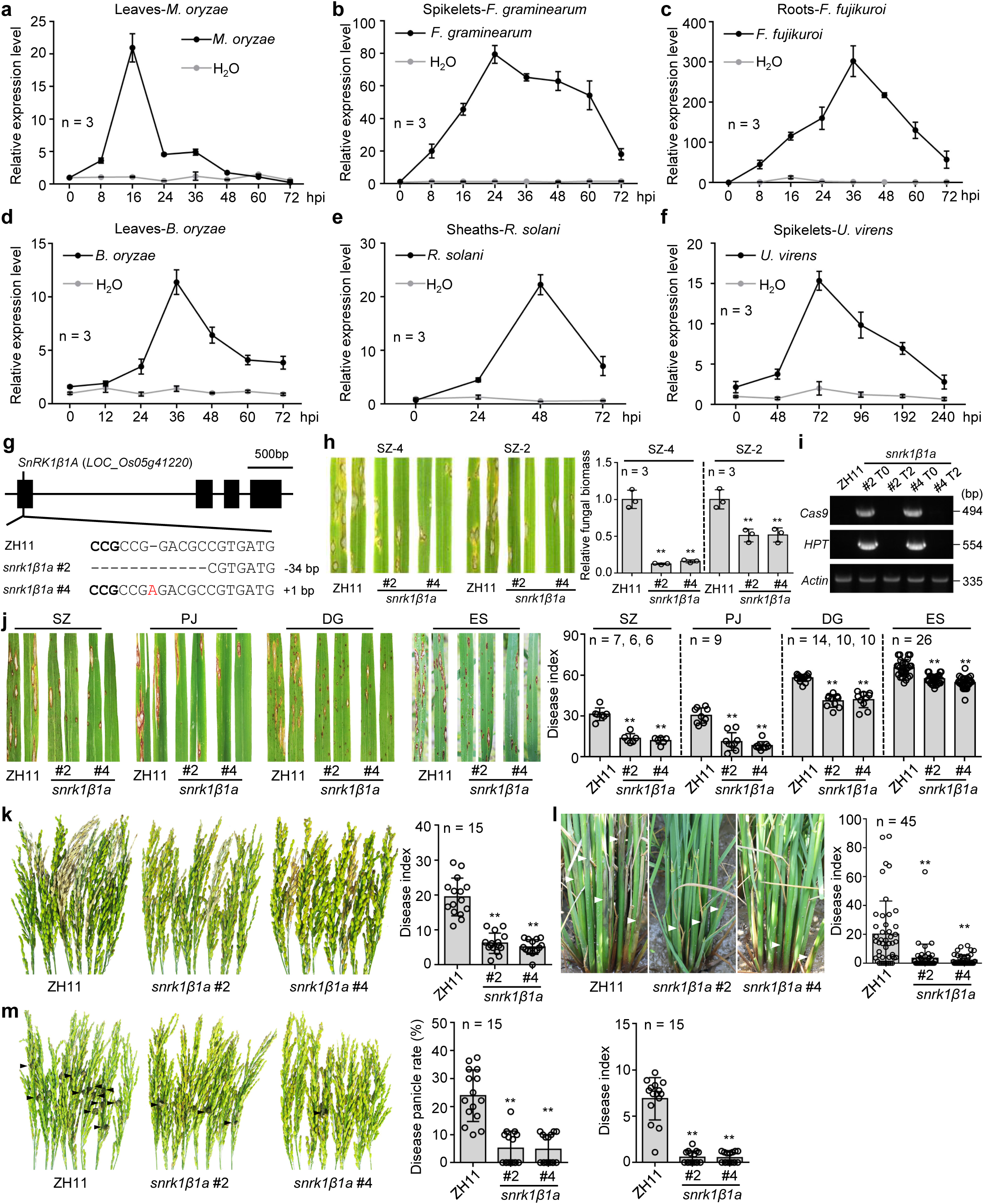
*SnRK1*β*1A* is induced upon infection by rice major fungal pathogens and its disruption enhances resistance to multiple fungal diseases in rice fields. **a-f,** Expression levels of *SnRK1*β*1A* in NPB after infection by *M. oryzae* (**a**), *F. graminearum* (**b**), *F. fujikuroi* (**c**), *B. oryzae* (**d**) *R. solani* (**e**) and *U. virens* (**f**) at indicated time points. H_2_O treatment served as a control. **g,** Schematic representation of the *SnRK1*β*1A* gene structure and gene editing sites. Bold letters indicate PAM, “-” indicates nucleotide deletion, red letter indicates nucleotide insertion. **h,** Infection assay showing that *snrk1*β*1a* mutants are resistant to diverse isolates of *M. oryzae*, including SZ-4 and SZ-2. Fungal biomass was measured by qPCR at 5 dpi. **i,** Identification of transgene-free *snrk1*β*1a* mutants. Primers specific to the *Cas9*, *HPT* and *Actin* genes, respectively, were used in genotyping. The *Actin* gene was used as the DNA quality control. **j,** *snrk1*β*1a* lines showed significantly reduced blast disease indices as compared with its WT ZH11 in rice fields of Shangzhuang (SZ), Panjin (PJ), Donggang (DG) and Enshi (ES). **k-m,** *snrk1*β*1a* lines are resistant to panicle blast, sheath blight and false smut disease in DG field. The white triangle in (**l**) indicates the symptoms of sheath blight disease and the white triangle in (**m**) indicates the symptoms of false smut ball. **j**-**m** were from natural infection with multiple duplicates. Two-tailed Student’s t-test were used in (**h,j-m**), **p < 0.01.

**Extended Data Fig. 5.**
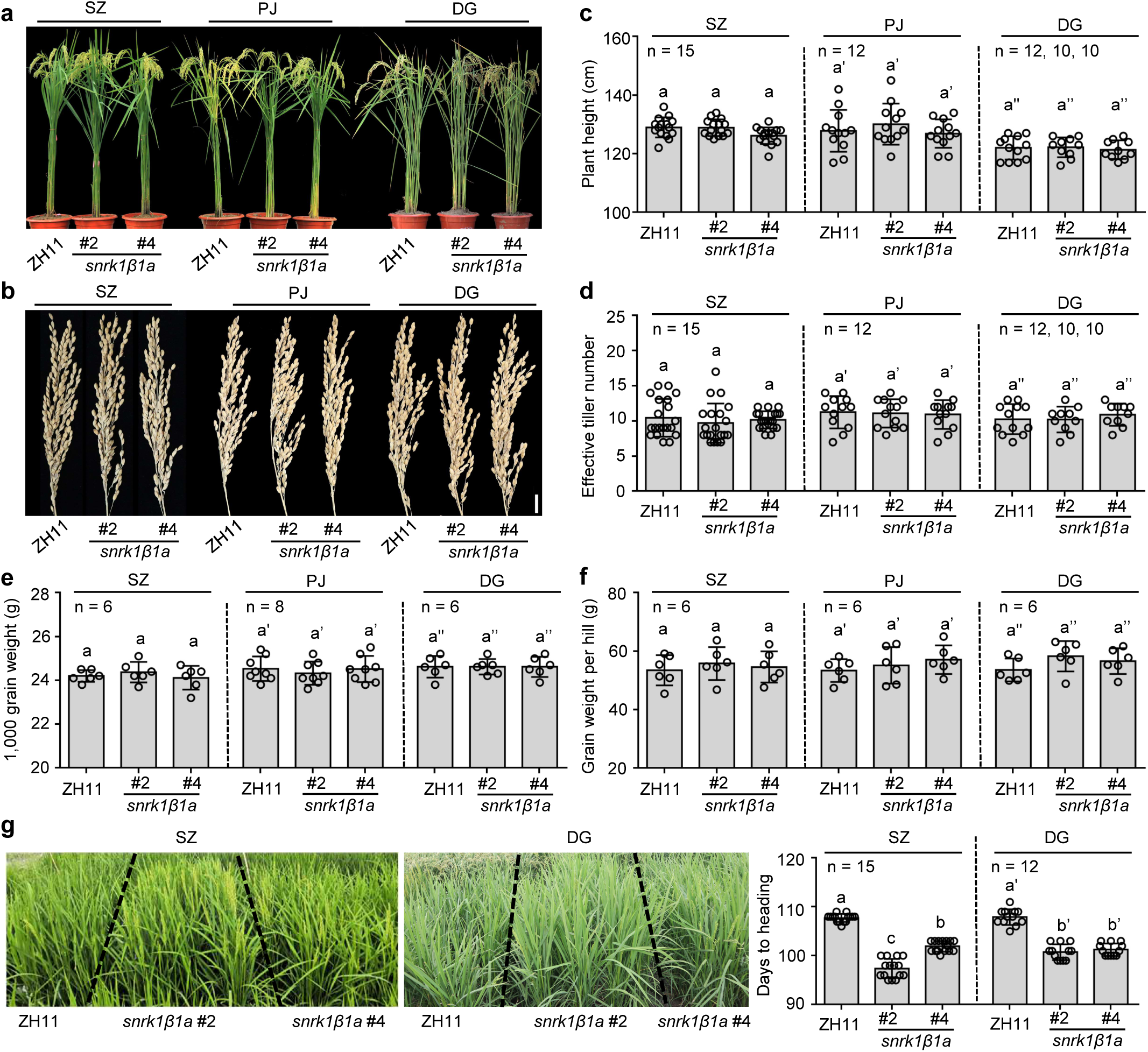
*snrk1*β*1a* mutants are similar to its WT in major agronomic traits except early heading. The plant morphology (**a**) and panicle morphology (**b**) of ZH11 and *snrk1*β*1a* in SZ, PJ, and DG fields. The main agronomic traits, including plant height (**c**), effective tiller number (**d**), 1,000 grains weight (**e**) and grain weight per hill (**f**) were investigated. **g**, *snrk1*β*1a* shows early heading. The rice lines were grown in the SZ and DG field and the heading time is counted. One-way ANOVA followed by post-hoc Turkey tests were used in (**c-g**); the Same letters in (**c-f**) indicate no significantly different groups, and the letters in (**g**) indicate significantly different groups.

**Extended Data Fig. 6.**
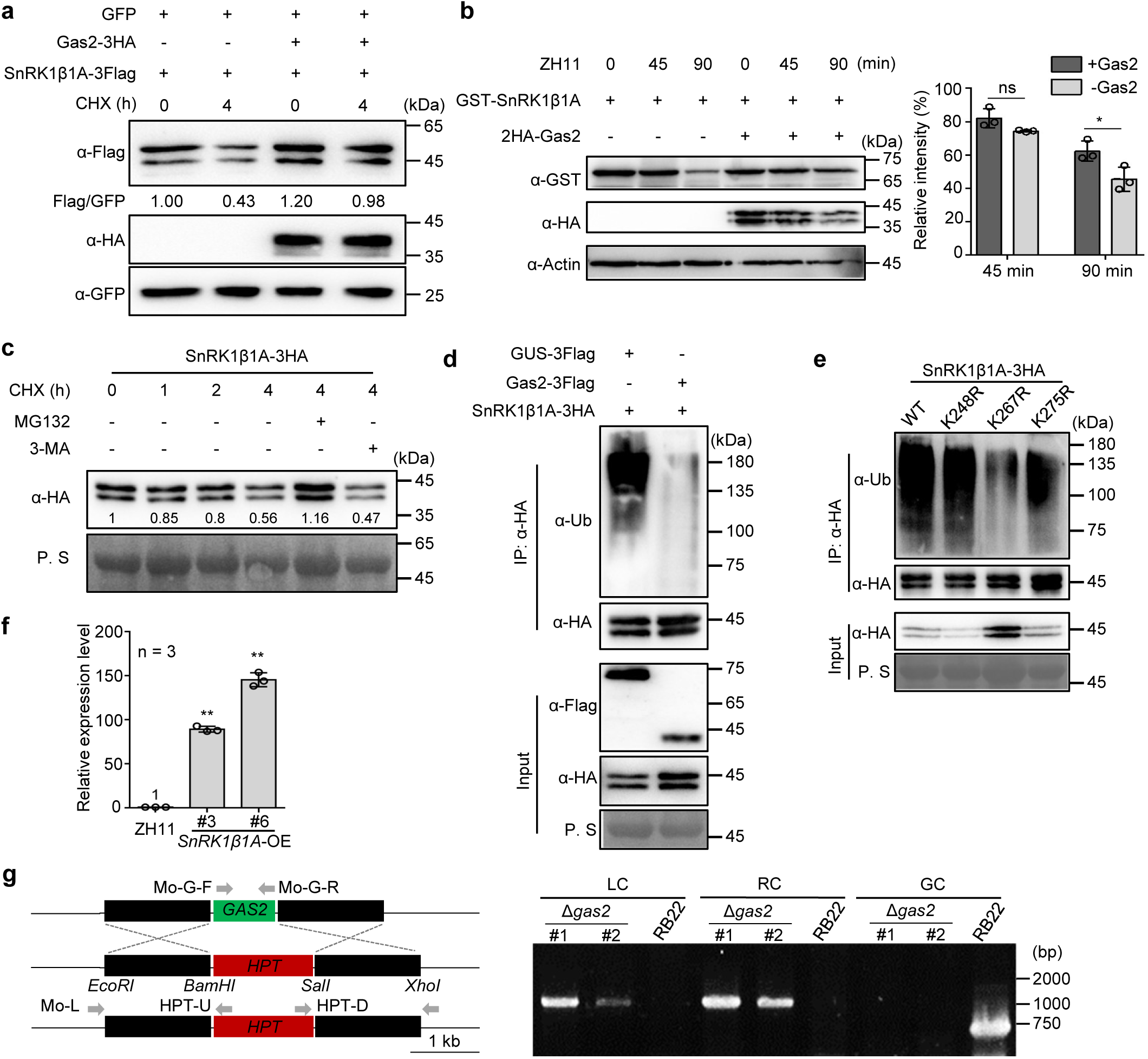
SnRK1β1A is degraded via the 26S proteasome system. **a,** SnRK1β1A-3Flag was more accumulated in rice protoplasts co-expressing Gas2-3HA and GFP, but not in those co-expressing GFP. WB was conducted using α-Flag, α-HA and α-GFP antibodies. Cycloheximide (CHX) was added in the protein extracts for 4 hours to inhibit protein synthesis. **b,** 2HA-Gas2 reduced degradation of purified GST-SnRK1β1A by total protein extracts from rice leaves. Proteins were detected within a 0-90 minute time course by WB using α-GST and α-HA antibodies, with α-actin as a loading control. **c,** The 26S proteosome inhibitor MG132 but not the autophagy inhibitor 3-Methyladenine (3-MA) suppressed degradation of SnRK1β1A-3HA. *N. benthamiana* leaves were injected with *SnRK1*β*1A-3HA* vector, and at 36 hpi, were injected with MG132/CHX or 3-MA/CHX. At 4 hours post the chemical injection, proteins were extracted for WB detection using anti-HA antibody, with P. S as a loading control. **d,e,** SnRK1β1A-3HA with Gas2-3Flag or GUS-3Flag (**d**), SnRK1β1A^K248R^-3HA, SnRK1β1A^K267R^-3HA and SnRK1β1A^K275R^-3HA (**e**) were transiently expressed in *N. benthamiana* leaves. The proteins were immunoprecipitated with α-HA beads for detecting their ubiquitination levels by WB using α-Ubiquitin antibody. **f,** RT-qPCR with *OsActin* as a control to confirm *SnRK1*β*1A* overexpression in transgenic ZH11 (*SnRK1*β*1A*-OE). Two-tailed Student’s t-tests were used, **p < 0.01. **g,** Δ*gas2* knockout mutants in RB22 isolate were generated by homologous recombination strategy (Top) and checked by PCR (bottom). Genomic DNA isolated from WT strain RB22 and the Δ*gas2* mutants was amplified by PCR using specific primers.

**Extended Data Fig. 7.**
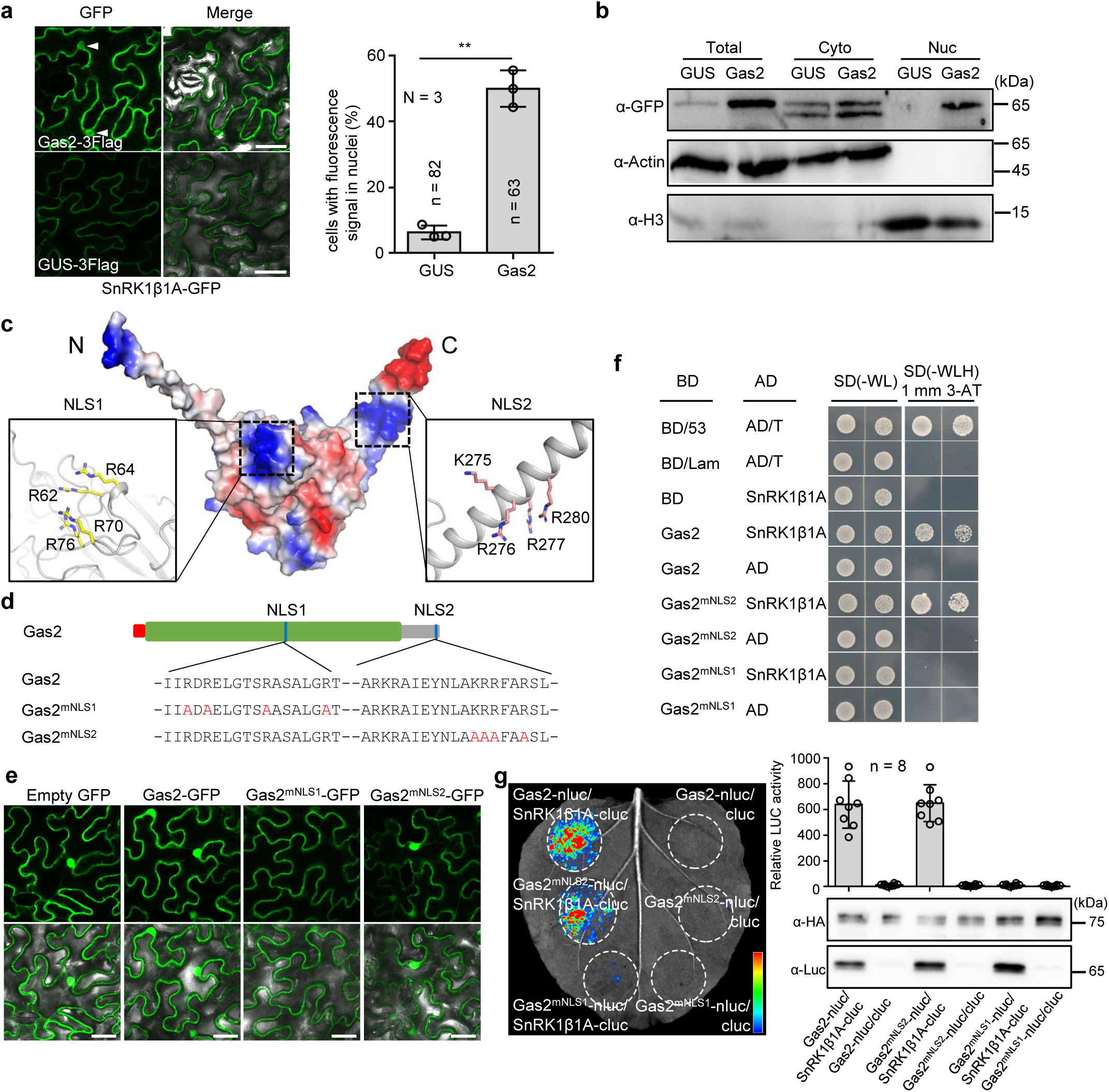
Gas2 leads SnRK1β1A into the nuclei of tobacco cells and has a functional nuclear localization signal. **a,** SnRK1β1A-GFP signals were more observed in the nucleus of *N. benthamiana* cells co-expressing Gas2-3Flag than those co-expressing GUS-3Flag. Cells with fluorescence signals in nuclei were counted. The white triangle indicates the nucleus. Scale bars, 25 μm. Two-tailed Student’s t-test were used, **p < 0.01. **b,** SnRK1β1A-GFP signals were more detected by the α-GFP antibody in the nucleus of *N. benthamiana* cells co-expressing Gas2-3Flag than those co-expressing GUS-3Flag. α-Actin and α-H3 served as the cytoplasmic and nuclear markers, respectively. **c,** Electrostatic surface potential of Gas2 predicted by PyMol. Positively charged and negatively charged surfaces of Gas2 are displayed in blue and red, respectively. The positively charged side chains of amino acids are indicated as putative nuclear localization signals (NLS). **d,** Mutation strategy of predicted NLSs in Gas2. Gas2^mNLS1^ and Gas2^mNLS2^ were obtained by mutating all lysine/arginine to alanine in NLS1 and NLS2, respectively. **e,** Subcellular localizations of Gas2-GFP and its NLS mutants in *N. benthamiana.* Green fluorescence was observed after 36 hours of infiltration using confocal microscopy. Scale bars, 25 μm. **f, g,** Y2H assays (**f**) and LCI assays (**g**) showed Gas2^mNLS1^ fails to interact with SnRK1β1A.The relative luciferase activity was measured, and protein expression was confirmed by immunoblot analysis with α-HA and α-Luc antibodies.

**Extended Data Fig. 8.**
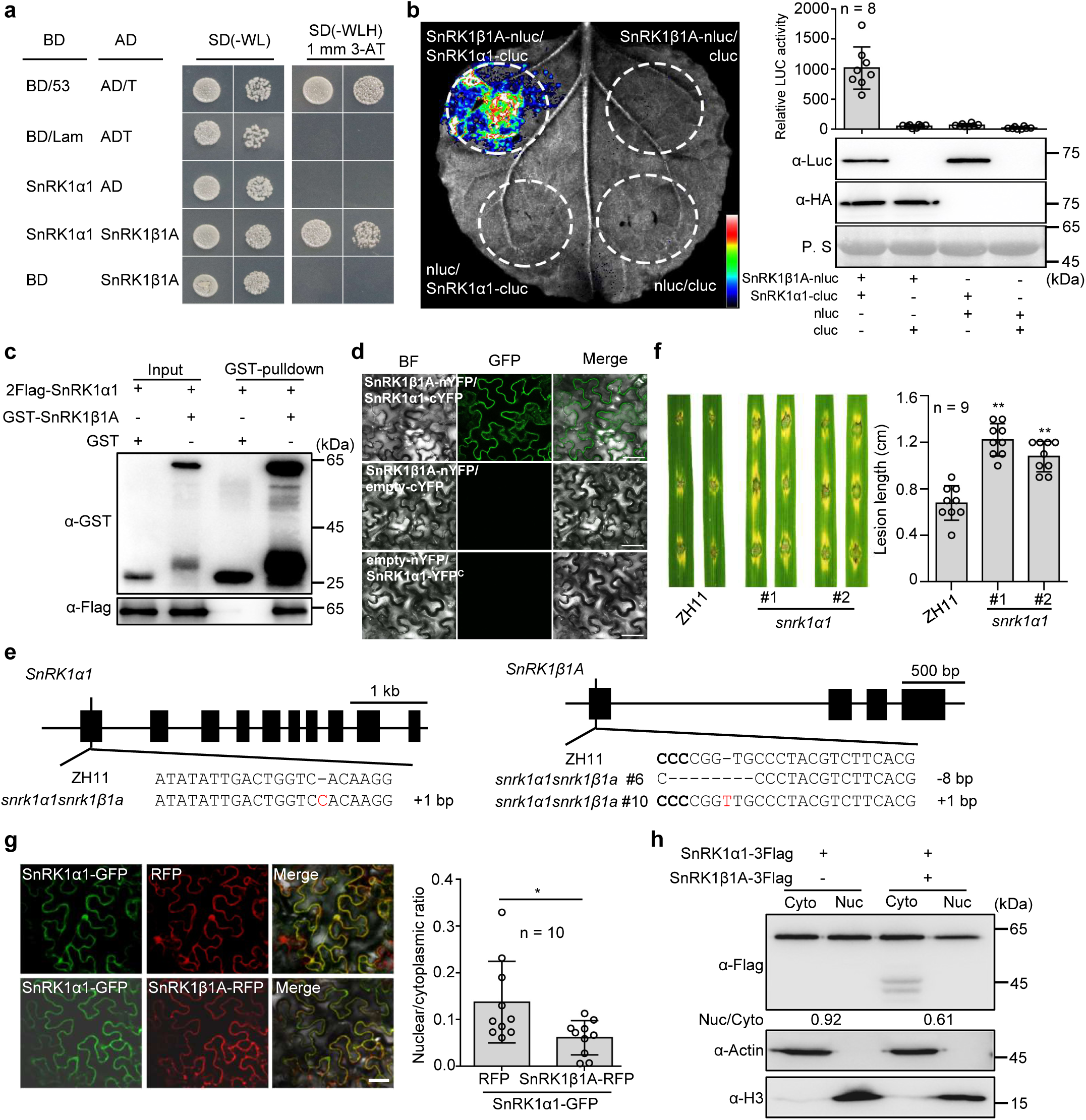
SnRK1β1A interacts with SnRK1α1 and inhibits nuclear localization of SnRK1α1. **a-d,** Interaction of SnRK1β1A with SnRK1α1 detected by Y2H assays (**a**), LCI assays (**b**), GST-pulldown assays (**c**) and BiFC assays (**d**). The relative luciferase activity was measured and protein expression was confirmed by immunoblot analysis with α-HA and α-Luc antibodies in (**b**). Scale bars in (**d**), 25 μm. **e,** Schematic representation of the *SnRK1*α*1* and *SnRK1*β*1A* structure and gene editing sites in the *snrk1*β*1asnrk1*α*1* double mutant. Bold letters indicate PAM, “-” indicates nucleotide deletion, and red letter indicates nucleotide insertion. **f,** Knockout of *SnRK1*α*1* increased susceptibility to blast. Rice leaves were punch-inoculated with *M. oryzae* isolate SZ-5. The lesion length was measured at 5 dpi. **g,** SnRK1β1A inhibited nuclear localization of SnRK1α1 in *N. benthamiana* cells. SnRK1α1-GFP co-expressed with SnRK1β1A-RFP or free RFP in *N. benthamiana*. Green fluorescence was observed after 36 hours of infiltration using confocal microscopy. Scale bar, 25 μm. **h,** WB showing less distribution of SnRK1α1-3Flag in the nucleus when SnRK1β1A-3Flag was co-expressed in *N. benthamiana* leaves. α-Flag antibody was used to detect the proteins in the cytoplasmic and nuclear fractionations with α-Actin and α-H3 as the cytoplasmic and nuclear markers, respectively. The nuclear to cytoplasmic ratio (**g,h**) was determined using ImageJ software. Two-tailed Student’s t-tests were used in **f,g**. *p < 0.05, **p < 0.01.

**Extended Data Fig. 9.**
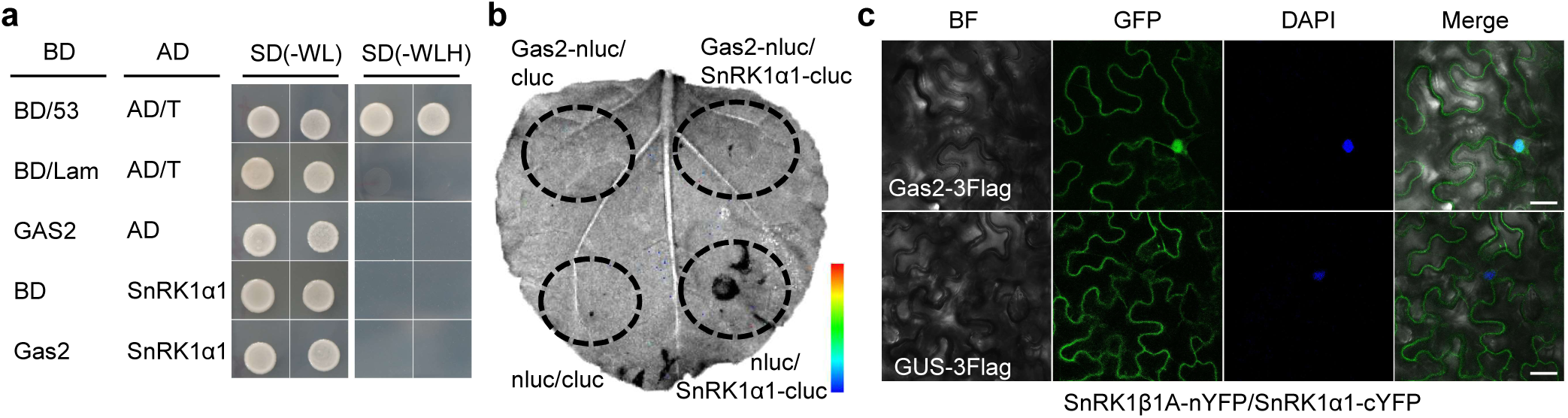
Gas2 is unable to interact with SnRK1α1 but enhances the interaction of SnRK1β1A with SnRK1α1 in the nucleus of plant cells. **a,** Y2H and **b,** LCI assays showing Gas2 unable to interact with SnRK1α1. **c,** BiFC assays showing the Gas2 enhanced interaction of SnRK1β1A with SnRK1α1 in the nucleus. Scale bars, 25 μm.

